# Optimal strategies for learning multi-ancestry polygenic scores vary across traits

**DOI:** 10.1101/2021.01.15.426781

**Authors:** B.C.L. Lehmann, M. Mackintosh, G. McVean, C.C. Holmes

**Affiliations:** Department of Statistical Science, University College London, UK; Genomics England, London, UK; Big Data Institute, University of Oxford, UK; Department of Statistics, University of Oxford, UK; The Alan Turing Institute, London, UK

## Abstract

Polygenic scores (PGSs) are individual-level measures that aggregate the genome-wide genetic predisposition to a given trait. As PGS have predominantly been developed using European-ancestry samples, trait prediction using such European ancestry-derived PGS is less accurate in non-European ancestry individuals. Although there has been recent progress in combining multiple PGS trained on distinct populations, the problem of how to maximize performance given a multiple-ancestry cohort is largely unexplored. Here, we investigate the effect of sample size and ancestry composition on PGS performance for fifteen traits in UK Biobank. For some traits, PGS estimated using a relatively small African-ancestry training set outperformed, on an African-ancestry test set, PGS estimated using a much larger European-ancestry only training set. We observe similar, but not identical, results when considering other minority-ancestry groups within UK Biobank. Our results emphasise the importance of targeted data collection from underrepresented groups in order to address existing disparities in PGS performance.

## Introduction

Polygenic scores (PGS) are composite, quantitative measures that aim to predict complex traits from genetic data. As well as providing insights into the genetic architecture of complex traits, PGS have considerable clinical potential for screening and prevention strategies [1, 2]. Largely driven by significant increases in sample sizes, the predictive utility of PGS has improved substantially in recent years for a variety of traits [3], including cardiovascular disease [4], breast cancer [5], and type I diabetes [6].

These improvements, however, have largely been limited to populations of European ancestry [7, 8, 9], reflecting the lack of diversity in genomic samples collected to date [9]. Moreover, predictive performance decreases with genetic distance from the training population [10, 11, 12]. The lack of transferability of PGS across ancestries may be due to a number of factors, including population differences in allele frequencies and linkage disequilibrium [9, 11]. There is also increasing evidence that the underlying variant effects differ across ancestries [13, 14], which may be due to gene-by-gene or gene-by-environment (GxE) interactions [15, 16]. In addition, GxE interactions for people of African ancestry may be different between those living in the UK and those living in South Africa, say. This lack of transferability raises one of the most important technical and ethical challenges in the clinical utility and applications of PGS due to their potential to exacerbate health inequalities [9].

There are major, ongoing initiatives to collect genomic data from traditionally under-represented groups, such H3Africa [17], that aim to address the lack of global genetic diversity in research data. However, it may take many years to collect sufficient data to reduce the disparities in PGS performance. Statistical methods may provide an alternative, short-term, cost-effective and complementary potential solution to mitigate against the negative effects of the lack of diversity in genomic datasets, by using modelling techniques to make use of all the existing data available, while allowing for some differences between groups.

There has been a growing interest in statistical methods to improve the transferability of PGS, which have thus far focused on GWAS-derived PGS, i.e. PGS based on summary statistics from a genome-wide association study. These integrate results from GWAS trained on distinct populations at different stages of the PGS estimation. For example, Grinde et al. [18] use European GWAS results to select variants, and then estimate the variant weights using the non-European GWAS results. Marquez-Luna et al. [19] propose a ‘multi-ethnic’ PGS by combining scores trained separately on different populations. Weissbrod et al. [20] expand on this approach by further leveraging functionally informed fine-mapping to estimate the PGS weights. In a related approach, Ruan et al. [21] incorporate linkage disequilibrium (LD) information to directly estimate effect sizes using GWAS results from two populations. To account for admixture, Cavazos & Witte [22] propose local ancestry weighting to construct individualised PGS. These efforts have yielded promising improvements in PGS performance, though there remains a significant gap in predictive accuracy between European and non-European target populations.

Here, we investigate the use of multiple-ancestry datasets, such as UK Biobank [23], to estimate PGS for a range of anthropometric, blood-sample, and disease traits, with the explicit objective of improving predictive accuracy for under-represented ancestries. Specifically, we ask whether there are consistent optimal strategies for borrowing information across ancestry groups to maximize prediction accuracy in groups that have small sample sizes in available resources. For each trait, we construct training sets with varying numbers of individuals from each ancestry to assess the effect of sample size and composition on PGS accuracy, using both simulated data and data from UK Biobank [23]. Moreover, and to counteract the imbalance of ancestries in a multiple-ancestry training set, we investigate the use of an importance reweighting approach that places more weight on underrepresented ancestries during training. Importance reweighting is a standard statistical technique used in survey sampling that aims to account for differences between the sampling population and the population of interest [24]. Given the availability of individual-level information in biobanks, we estimate PGS using regularised regression applied to full genotype and covariate data (as opposed to genome-wide association summary statistics) to avoid introducing additional artefacts into our analysis via the reliance on assumptions about genotype and covariate correlation structure (including linkage disequilibrium) and GWAS methodology.

Our results show that the impact of sample size and composition on predictive performance is highly variable across traits. For some traits, polygenic scores estimated using a relatively small number of minority-ancestry individuals outperformed on a minority-ancestry test set scores estimated using a much larger number of European-ancestry individuals. Moreover, adding European-ancestry individuals to the training set did not always improve performance and in some cases even led to poorer performance. Although importance weighting yields moderate improvement in performance for some traits, we find that sample size is a much more prominent factor, highlighting the limitations of statistical corrections and the importance of collecting more data from a more diverse range of participants.

## Results

### Overview of methods

To investigate the effect of sample size and ancestry composition on polygenic score performance, we used both simulated data and real data from UK Biobank [23]. We initially focus on PGS strategy for African-ancestry groups, given that predictive accuracy using European-ancestry PGS is consistently worst out of all the major ancestry groups [9, 12]. Simulated data was generated using the simulation engine `msprime` [25] in the standard library of population genetic simulation models `stdpopsim` v0.1.2 [26] to generate African-ancestry and European-ancestry genotypes. We also considered a range of quantitative and binary traits from UK Biobank, using imputed genotypes along with inferred genetic ancestry labels made available by the Pan-UKBB initiative [27].

For both the simulated and real data, we constructed a range of training sets controlling the number of European-ancestry and minority-ancestry individuals. We considered three types of training sets: a single-ancestry set consisting only of European-ancestry individuals, a single-ancestry set consisting of minority-ancestry individuals, and a dual-ancestry set consisting of both European-ancestry and minority-ancestry individuals. We denote a PGS trained on a XYZ-ancestry training set as PGS_XYZ_-for example PGS_EUR_. Similarly, we refer to a PGS trained on a dual-ancestry (respectively minority-ancestry) training set as PGS_dual_ (respectively PGS_min_).

For each training set, we estimated PGS using L1-regularised regression, also known as the LASSO [28], which has previously been used in the context of genetic risk prediction (see for example [29, 30, 31, 32]). To account for the imbalance in sample size numbers in the dual-ancestry training sets, we also estimated PGS using an importance reweighted LASSO, upweighting the non-European-ancestry individuals. Following Martin et al. [9], we assessed the predictive performance of a PGS using partial *r*^2^ relative to a covariate-only model. See Figure 1 for a schematic diagram of the methods used and Materials and Methods for full details.

**Figure 1:**
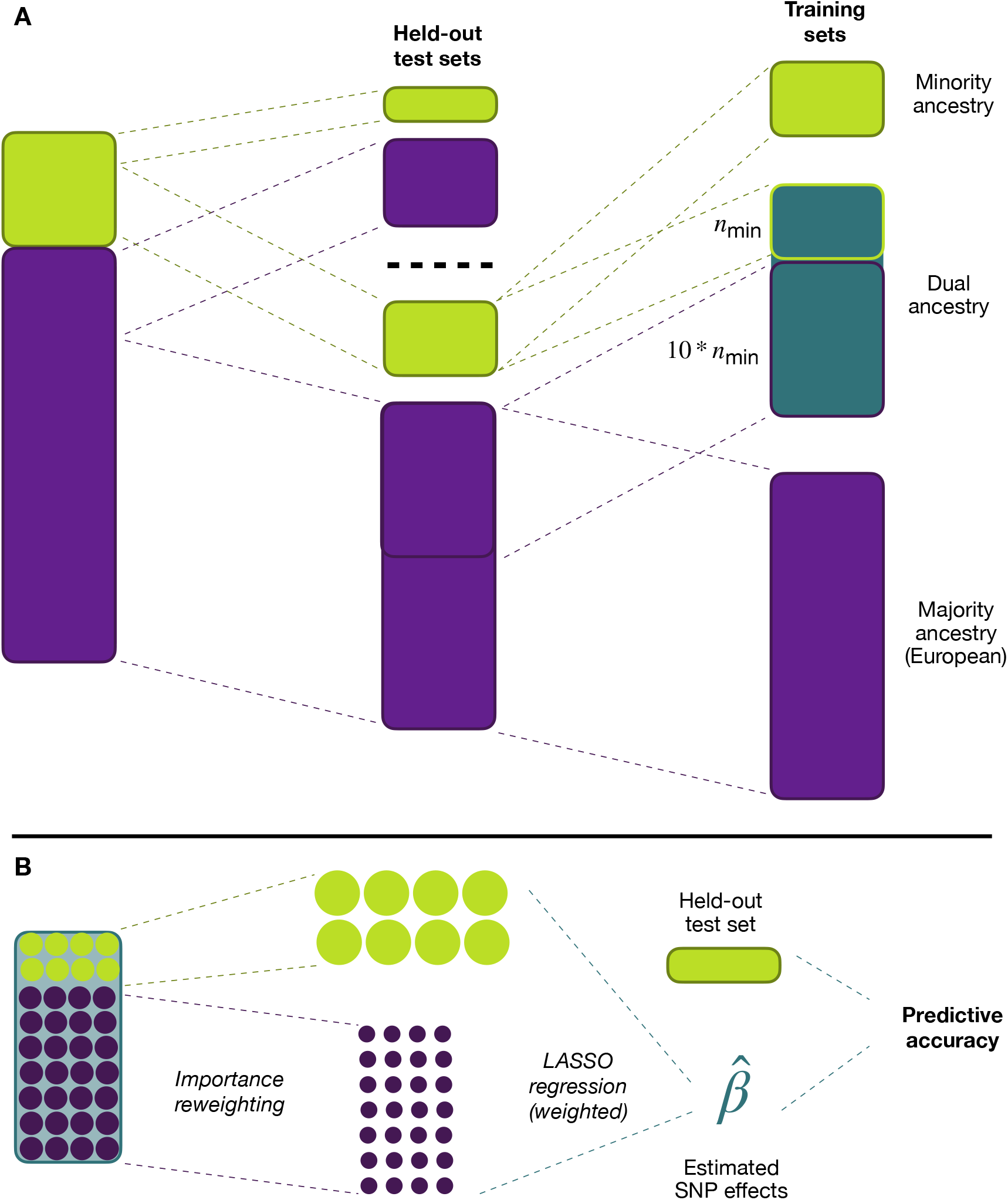
Overview of methods. A) To evaluate the different PGSs, we performed various splits of the available data. Firstly, we held out test sets of 20% of individuals in each ancestry group. From the remaining 80%, we constructed three types of training sets: a single-ancestry set consisting only of European-ancestry individuals, a single-ancestry set consisting of non-European-ancestry individuals, and a dual-ancestry set consisting of both European-ancestry and non-European-ancestry individuals. For each training set, we used another 20% of the data to select the regularisation parameter in the LASSO. B) For the dual-ancestry training set, we used an importance weighted LASSO, assigning higher weights to individuals in the minority-ancestry group. See Materials and Methods for full details.

### Single-ancestry PGS can outperform dual-ancestry PGS despite being trained on fewer individuals

We first set out to evaluate the relative performance of single-ancestry PGS versus dual-ancestry PGS through simulation. An important factor in the lack of transferability of PGS across ancestries is the difference in causal effect sizes [11], with the correlation of causal effect sizes between ancestry groups, *ρ*, has been estimated to be significantly less than one across a range of common traits [14, 33]. To assess the impact of differences in causal effect sizes on the relative predictive performance of PGS strategies, we varied the trans-ancestry causal effect correlation, *ρ* = 0.5, 0.6,…, 0.9, generating 10 quantitative traits for each value. We assume that all causal variants are genotyped, thus avoiding any differences in PGS accuracy that may arise from imperfect tagging [11]. We note that in practice imputation does not fully address the issue of imperfect tagging, due to differences in imputation quality across ancestry groups [7, 9].

For each trait we created five African-ancestry training sets with *n_AFR_* = 2 000, 4 500, 7 714, 12 000, 18 000 African-ancestry individuals. We also created five dual-ancestry training sets made up of the corresponding African-ancestry training set supplemented with *n_EUR_* = 18000 European-ancestry individuals. To obtain PGS from the African-ancestry training sets, we used unweighted LASSO regression, while for the dual-ancestry training sets, we used the importance reweighted LASSO with *γ* = 0, 0.1,…, 1. The case *γ* = 0 corresponds to no reweighting, that is, the standard LASSO. The case *γ* =1 corresponds to inverse proportion reweighting, so that in total, African-ancestry individuals and European-ancestry individuals have equal weight. Note that the African-ancestry training set is equivalent to the limiting case *γ* → ∞ whereby zero weight is placed on European-ancestry individuals. We quantified predictive performance of a PGS in terms of the *predictive gap*: the difference between variance explained by the PGS, *r*^2^, and SNP heritability *h*^2^ (the maximal variance explained by any PGS). See Simulation Study for full details.

For both PGS_AFR_ and PGS_dual_, the predictive gap for African-ancestry individuals decreased substantially as the number of African-ancestry individuals in the training sets increased (Figure 2). For example, when *n_AFR_* = 2 000 and the correlation between genetic effects ρ was 0.7, the mean predictive gap of the unweighted PGS_dual_ (*γ* = 0) was just over 0.2 for the African-ancestry test sets, compared to 0.03 for the European-ancestry test sets. While the performance on Europeans did not change markedly as the number of African-ancestry training set individuals increased, the African-ancestry predictive gap fell to approximately 0.11 and 0.06 when *n_AFR_* = 7714 and *n_AFR_* = 18 000 respectively. As expected, the discrepancy between the two ancestry groups decreased as the correlation ρ between the ancestry-specific genetic effects increased.

**Figure 2:**
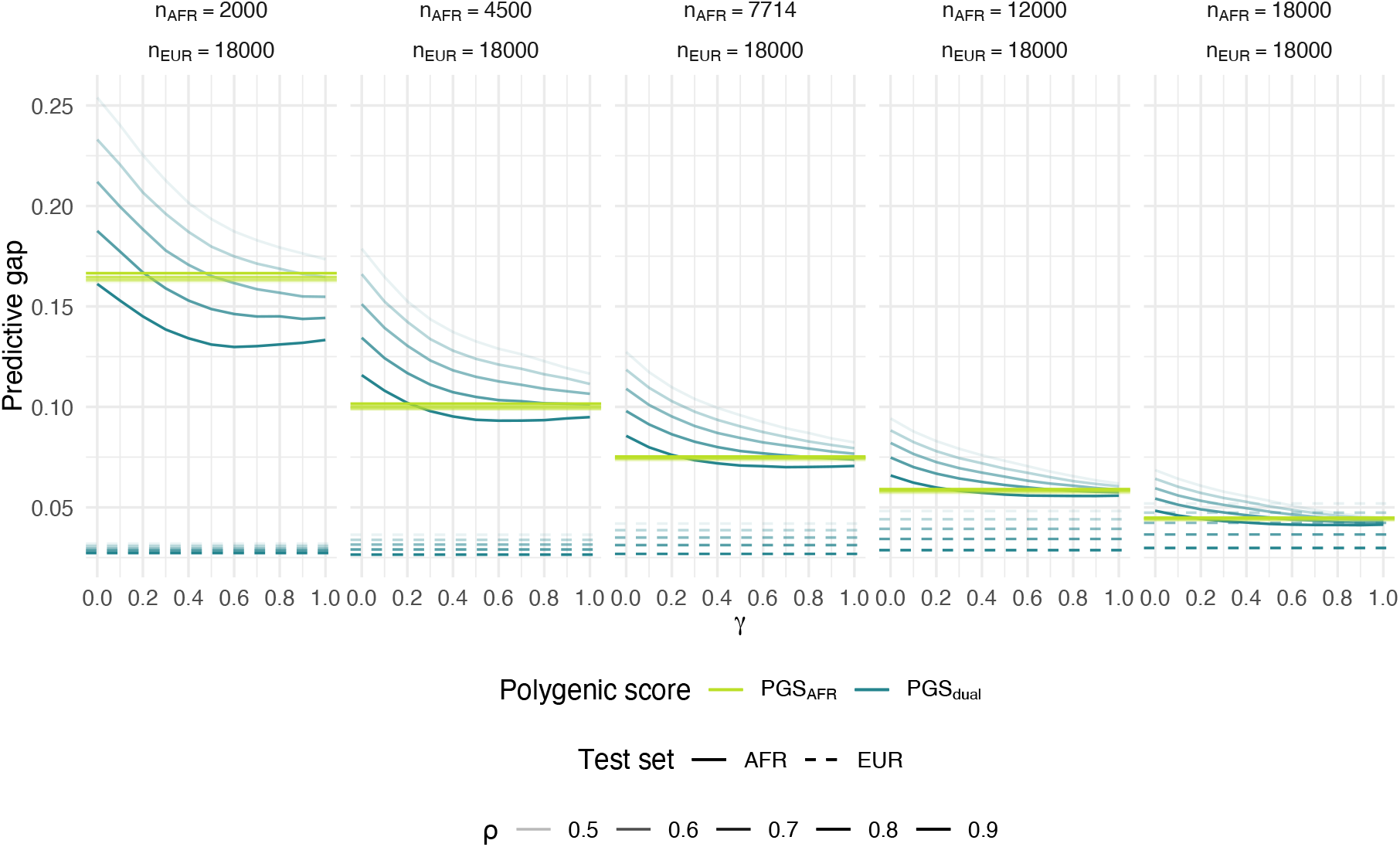
Simulation study: predictive gap against number of African-ancestry individuals in training set. Each panel corresponds to a different number of African-ancestry training set individuals from *n_AFR_* = 2000 to *n_AFR_* = 18000. The training sets for PGS_dual_ (blue lines) consisted of the corresponding African-ancestry training set for PGS_AFR_ (yellow lines), along with *n_EUR_* = 18 000 European-ancestry individuals. Each line represents the mean predictive gap across 50 repetitions. The horizontal dashed lines correspond to the predictive gap for European-ancestry test sets based on an unweighted LASSO, while the solid lines correspond to the predictive gap for African-ancestry test sets. The correlation of genetic effects between ancestries *ρ* was varied from 0.5 (lighter lines) to 0.9 (darker lines).

In the absence of reweighting (*γ* = 0), PGS_AFR_ generally outperformed PGS_dual_, despite the latter being trained on more individuals - the dual-ancestry training sets consist of the African-ancestry training sets along with 18000 European-ancestry individuals. This was particularly evident when the correlation between ancestry-specific genetic effects was lower (*ρ* = 0.5).

Importance reweighting generally had a positive impact on predictive performance, reducing the difference between PGS_AFR_ and PGS_dual_. When the number of African-ancestry training individuals was low (*n_AFR_* = 2000), the importance-reweighted PGS_dual_ outperformed PGS_AFR_ when the correlation between ancestry-specific genetic effects was relatively high (*ρ* ≥ 0.7).

The effect on predictive performance depended on the degree of reweighting, quantified by *γ*. As *γ* increased from 0 to 1, the predictive gap typically decreased. When the correlation between ancestry-specific genetic effects was high (ρ = 0.9) and the number of African-ancestry samples low (*n_AFR_* = 2000), the predictive gap increased again for larger values of *γ*. This reflects a crucial trade-off of importance reweighting in this context: while the bias of African-ancestry genetic effect estimates may be lower with more reweighting, the increased variance of the weights results in a lower *effective* sample size. The effect of reweighting was relatively small compared to the effect of increased sample size. For *n_AFR_* = 2000 and *ρ* = 0.7, reweighting reduced the predictive gap from 0.21 to 0.15. The same improvement for the unweighted PGS_dual_ was seen when increasing the number of African-ancestry individuals to *n_AFR_* = 4 000, though we note that the PGS_AFR_ with *n_AFR_* = 4000 had an even lower predictive gap of 0.10. The amount of improvement with reweighting decreased only slightly as *n_AFR_* increased, and decreased more markedly as the correlation between genetic effects ρ increased.

This simulation study illustrates the relative performance of PGS_AFR_ and PGS_dual_ as a function of i) the number of minority-ancestry individuals in the respective training sets and ii) the correlation of genetic effects between ancestries. For both PGS_AFR_ and PGS_dual_, the between-ancestry disparity in predictive performance decreased as the number of minority-ancestry individuals in the training set increased. Despite being trained on fewer individuals, the PGS_AFR_ tended to outperform PGS_dual_, particularly when the correlation between ancestry-specific genetic effects was low. Importance reweighting generally reduced the predictive gap of PGS_dual_, especially when the sample size imbalance between ancestries was high. As this imbalance decreased, however, the reweighted PGS_dual_ did not outperform PGS_AFR_.

### Adding individuals from one ancestry does not always improve PGS performance for a different ancestry

Next, we sought to examine whether prediction performance in an underrepresented ancestry group can be boosted by including individuals from a different ancestry among the training samples. To do so, we used data from UK Biobank, estimating a range of PGS based on varying numbers of training individuals for a variety of traits (Table 1; see Materials and Methods for a description of how the traits were selected). In this analysis we focused on African-ancestry and European-ancestry individuals, where genetic ancestry labels were taken from Pan-UKBB.

**Table 1:** Number of individuals with non-missing trait value in UK Biobank by genetic ancestry group. SAIGE heritability estimates [70] were obtained from the Pan-UKBB initiative website [27].

| Ancestry | Trait | Total | Cases | $h^2$ |
| --- | --- | --- | --- | --- |
| AFR | Height | 6178 |  | 0.4603 |
| AMR | Height | 981 |  |  |
| CSA | Height | 8082 |  | 0.5709 |
| EAS | Height | 2689 |  | 0.4415 |
| EUR | Height | 361699 |  |  |
| MID | Height | 1543 |  |  |
| AFR | Mean corpuscular volume (MCV) | 5912 |  | 0.4465 |
| AMR | Mean corpuscular volume (MCV) | 958 |  | 0.8142 |
| CSA | Mean corpuscular volume (MCV) | 7956 |  | 0.3716 |
| EAS | Mean corpuscular volume (MCV) | 2622 |  | 0.2414 |
| EUR | Mean corpuscular volume (MCV) | 351672 |  | 0.1215 |
| MID | Mean corpuscular volume (MCV) | 1513 |  | 0.7393 |
| AFR | Asthma | 6257 | 741 | 0.0255 |
| AMR | Asthma | 989 | 106 | 0.0610 |
| CSA | Asthma | 8285 | 1017 | 0.0340 |
| EAS | Asthma | 2700 | 236 | 0.0170 |
| EUR | Asthma | 362511 | 42248 | 0.0748 |
| MID | Asthma | 1567 | 174 | 0.0301 |
| AFR | Female genital prolapse (FGP) | 3665 | 93 | 0.1222 |
| AMR | Female genital prolapse (FGP) | 641 | 27 |  |
| CSA | Female genital prolapse (FGP) | 3793 | 170 |  |
| EAS | Female genital prolapse (FGP) | 1786 | 31 |  |
| EUR | Female genital prolapse (FGP) | 194734 | 11043 | 0.0675 |
| MID | Female genital prolapse (FGP) | 662 | 33 |  |
| AFR | Body mass index (BMI) | 6168 |  | 0.3838 |
| EUR | Body mass index (BMI) | 361306 |  | 0.1090 |
| AFR | Eosinophill percentage | 5893 |  | 0.0739 |
| EUR | Eosinophill percentage | 351038 |  | 0.0958 |
| AFR | Erythrocyte distribution width | 5912 |  | 0.1331 |
| EUR | Erythrocyte distribution width | 351672 |  | 0.0936 |
| AFR | Lymphocyte count | 5893 |  | 0.1896 |
| EUR | Lymphocyte count | 351033 |  | 0.0962 |
| AFR | Monocyte count | 5893 |  | 0.2451 |
| EUR | Monocyte count | 351033 |  | 0.1105 |
| AFR | Mean platelet volume (MPV) | 5912 |  | 0.3315 |
| EUR | Mean platelet volume (MPV) | 351669 |  | 0.2062 |
| AFR | Platelet crit | 5912 |  | 0.2090 |
| EUR | Platelet crit | 351670 |  | 0.1298 |
| AFR | High light scatter reticulocyte count | 5748 |  | 0.3758 |
| EUR | High light scatter reticulocyte count | 345862 |  | 0.0967 |
| AFR | Atrial fibrillation (AFib) | 6257 | 125 | 0.0797 |
| EUR | Atrial fibrillation (AFib) | 362511 | 16476 | 0.0880 |
| AFR | Diverticular disease of the intestine (DDI) | 6257 | 285 | 0.0247 |
| EUR | Diverticular disease of the intestine (DDI) | 362511 | 30413 | 0.0615 |
| AFR | Hypothyroidism | 6257 | 124 | 0.0308 |
| EUR | Hypothyroidism | 362511 | 17496 | 0.1243 |

First, we held the number of African-ancestry individuals fixed at approximately 5000 (or 2 500 for female genital prolapse (FGP)), corresponding to 80% of African-ancestry individuals with non-missing data for the trait in question. We then constructed six training sets, varying the number of European-ancestry individuals from 0 to just under 50 000, so that the proportion of European-ancestry individuals in the training set ranged from 0% to 90%. For each training set, we used the unweighted LASSO to generate polygenic scores.

Predictive performance in terms of variance explained on European-ancestry individuals improved for each trait as the number of European-ancestry individuals in the training set increased (Figure 3). In contrast, predictive performance on African-ancestry individuals only increased markedly for height and mean platelet volume (MPV), with the improvement for MPV appearing to tail off between the two largest training sets. For two traits (mean corpuscular volume, reticulocyte count), predictive performance worsened as European-ancestry individuals were added to the training set. For the remainder of the traits, predictive performance stayed mostly constant.

**Figure 3:**
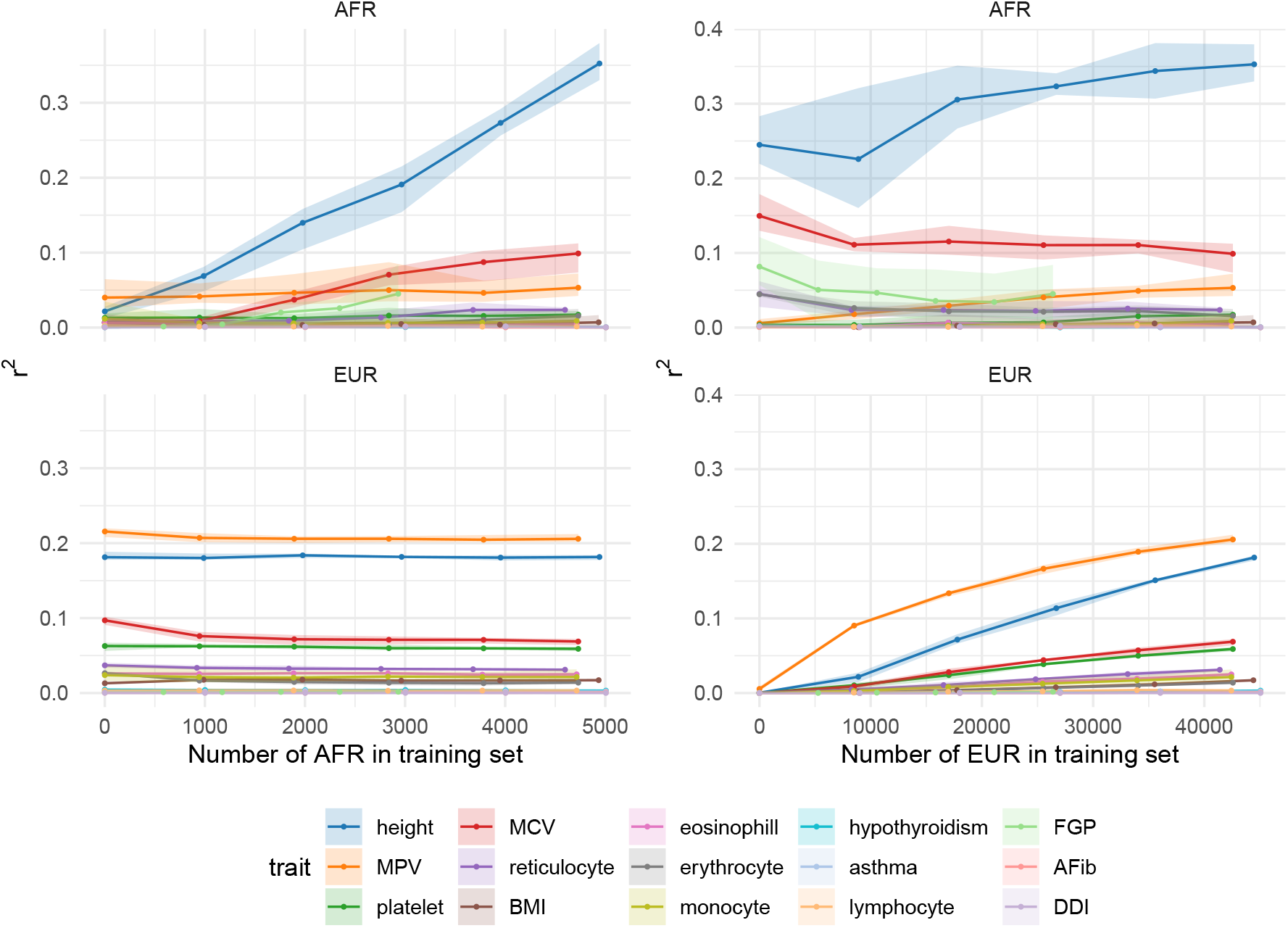
Partial *r*^2^ against sample size for 15 traits in UK Biobank. Left: We fixed the number of African-ancestry individuals in the training set at approximately 5 000 (2 700 for FGP) for each trait and varied the number of European-ancestry individuals so that the proportion of European-ancestry individuals in the training set ranged from 0% to 90%. While predictive performance on European-ancestry individuals increased monotonically with sample size for each trait (right panel), the effect on performance on African-ancestry individuals varied by trait (left panel). Right: Here, we instead fixed the number of European-ancestry individuals in the training set at approximately 50000 (26 000 for FGP) and varied the number of African-ancestry individuals from 0 to approximately 5 000 (2 700). The predictive performance on African-ancestry individuals increased for five traits:height, MCV, FGP, reticulocyte count, and platelet crit; and stayed largely stable for the remainder. Predictive performance on European-ancestry individuals remained stable for each of the fifteen traits. Error bars correspond to the range across five cross-validation rounds.

Next, we performed complementary analyses in which we held the number of European-ancestry individuals fixed at approximately 50 000, and instead varied the number of African-ancestry individuals from 0 to approximately 5 000. The predictive performance on African-ancestry individuals increased for five traits: height, mean corpuscular volume (MCV), FGP, high light scatter reticulocyte count (reticulocyte), and platelet crit (platelet). For the remainder of the traits, predictive performance stayed largely stable. Predictive performance on European-ancestry individuals also remained largely constant for each of the fifteen traits. These results demonstrate that the potential for increasing prediction performance by including samples from a different population can be limited. They also highlight substantial between-trait heterogeneity in response, suggestive of major differences in genetic architecture.

### Optimal ancestry composition of training sets varies among traits in UK Biobank

Next, we set out to assess the relative performance of single-versus dual-ancestry PGS in empirical data by considering the UK Biobank traits analysed in the previous section. We first considered three training sets: i) a European-ancestry training set of approximately 300 000 European-ancestry individuals, ii) a African-ancestry training set of approximately 5 000 African-ancestry individuals, and iii) a dual-ancestry training set made up of the African-ancestry training set combined with European-ancestry individuals so that the proportion of African-ancestry individuals was 10%. For the latter, we opted for this 90-10 split - thus not using all the available European-ancestry individuals - following the previous analysis that indicated the limited benefit to African-ancestry predictive performance of adding European-ancestry individuals to the training set. A secondary motivation was to limit the imbalance between the two ancestries in order to evaluate the effect of importance weighting. We used the importance weighted LASSO with *γ* = 0, 0.2,…, 1 to construct PGS_dual_, while for the single-ancestry training sets we just considered the standard, unweighted LASSO.

Figure 4 illustrates the predictive performance of the three above PGS for the 15 UK Biobank traits. In terms of African-ancestry predictive utility, PGS_AFR_ outperformed PGS_EUR_ on four traits - platelet crit (platelet), MCV, erythrocyte distribution width (erythrocyte), and FGP - despite the former sample size being orders of magnitude smaller (*n* ≈ 5 000 versus *n* ≈ 300 000). For each of these traits, the unweighted PGS_dual_ performed the same as or slightly worse than PGS_AFR_. While importance reweighting yielded a moderate improvement, the reweighted PGS_dual_ did not outperform PGS_AFR_, consistent with the findings from our simulation study.

**Figure 4:**
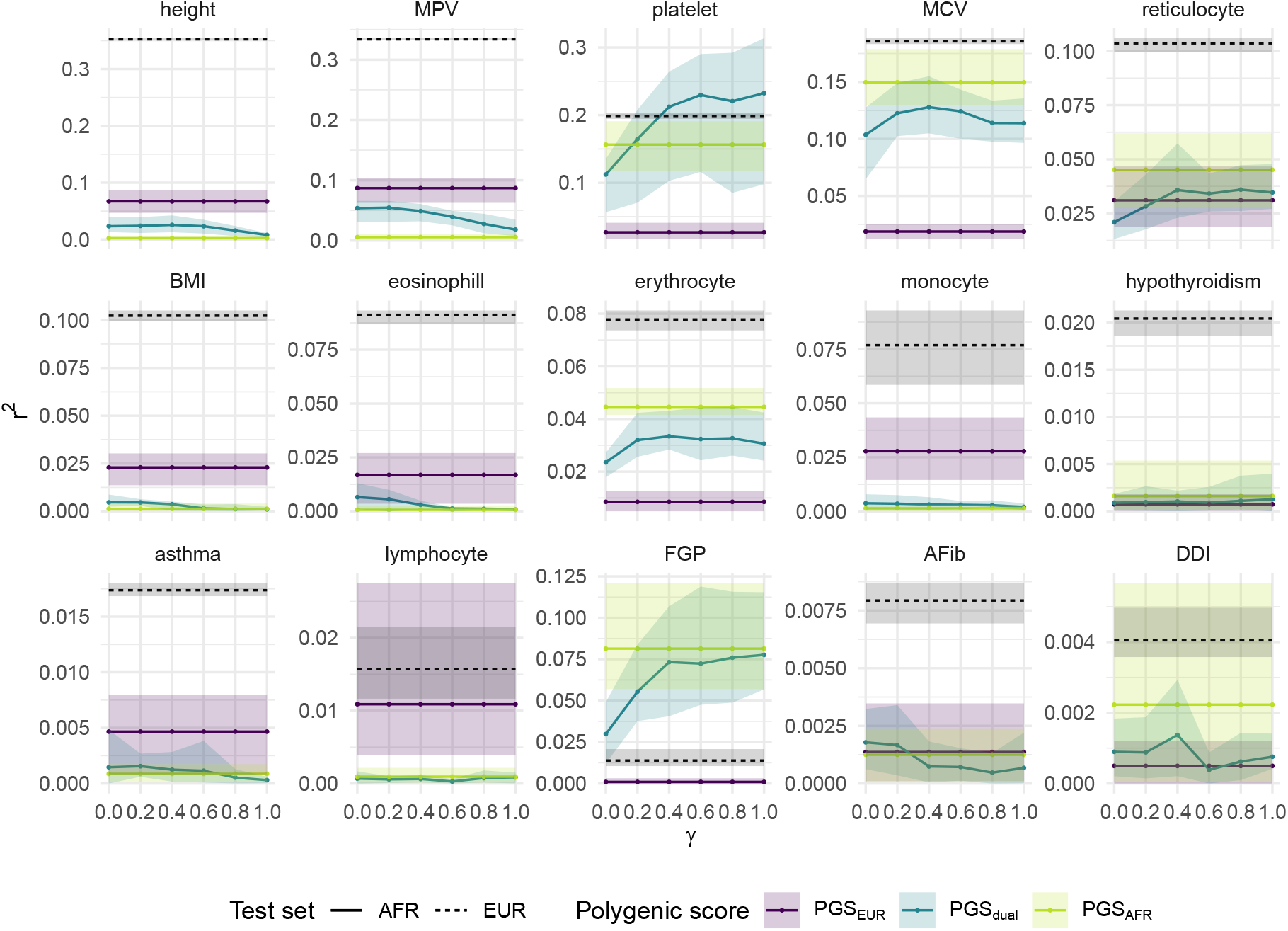
Partial *r*^2^ for PGS_EUR_, PGS_dual_, and PGS_AFR_ on 15 traits in UK Biobank. Predictive performance on an African-ancestry test set is shown by the solid lines. The dashed lines correspond to predictive performance on a European-ancestry test set using PGS_EUR_. The single-ancestry scores were estimated using a standard, unweighted LASSO. The dual-ancestry scores were constructed using an importance weighted LASSO with various degrees of reweighting *γ*. Traits are ordered according to partial *r*^2^ of PGS_EUR_ on the European-ancestry test set (note the varying y-axes). Error bars correspond to the range across five cross-validation rounds.

For four other traits - high light scatter reticulocyte count (reticulocyte), hypothyroidism, atrial fibrillation (AFib), and diverticular disease of intestine (DDI) - the predictive performance of all three PGS was largely similar. For the remaining traits - height, MPV, body mass index (BMI), eosinophill percentage, monocyte count, asthma and lymphocyte count - PGS_EUR_ performed best. We note that, with the exception of FGP, predictive accuracy of PGS_EUR_ for European-ancestry individuals was the same or higher than any of the PGS for African-ancestry individuals, consistent with previous investigations into the between-ancestry disparities of polygenic scores [7, 9].

To explore whether the variability in optimal PGS strategy extended to other subpopulations, we focused in on four of the above traits (height, MPV, asthma, and FGP) and ran the above analyses using different minority-ancestry groups: Admixed American ancestry (AMR), Central/South Asian ancestry (CSA), East Asian ancestry (EAS), and Middle Eastern ancestry (MID). Analyses were not run on FGP for AMR, EAS and MID ancestry groups as the number of cases was fewer than 50 in each group.

Figure 5 shows the predictive performance on the four traits across the five minority ancestry groups. For both height and asthma, PGS_EUR_had the best predictive performance for each of ancestry groups, with partial *r*^2^ for height varying from *r*^2^ = 0.08 for the African-ancestry group to *r*^2^ = 0.25 for the Admixed American ancestry group. On the other hand, for female genital prolapse, the PGS_min_ and the reweighted PGS_dual_ far outperformed PGS_EUR_, which offered almost no predictive utility. The picture for mean corpuscular volume was less consistent across ancestries: PGS_EUR_ was best for the Admixed American, Central/South Asian and Middle Eastern ancestry groups, while the PGS_min_ was best for the African-ancestry group. For the East Asian ancestry group, each of the three PGS had similar predictive performance. Results were qualitatively identical when using only genotyped SNPs instead of the full imputed sequence (SI Figure 1).

**Figure 5:**
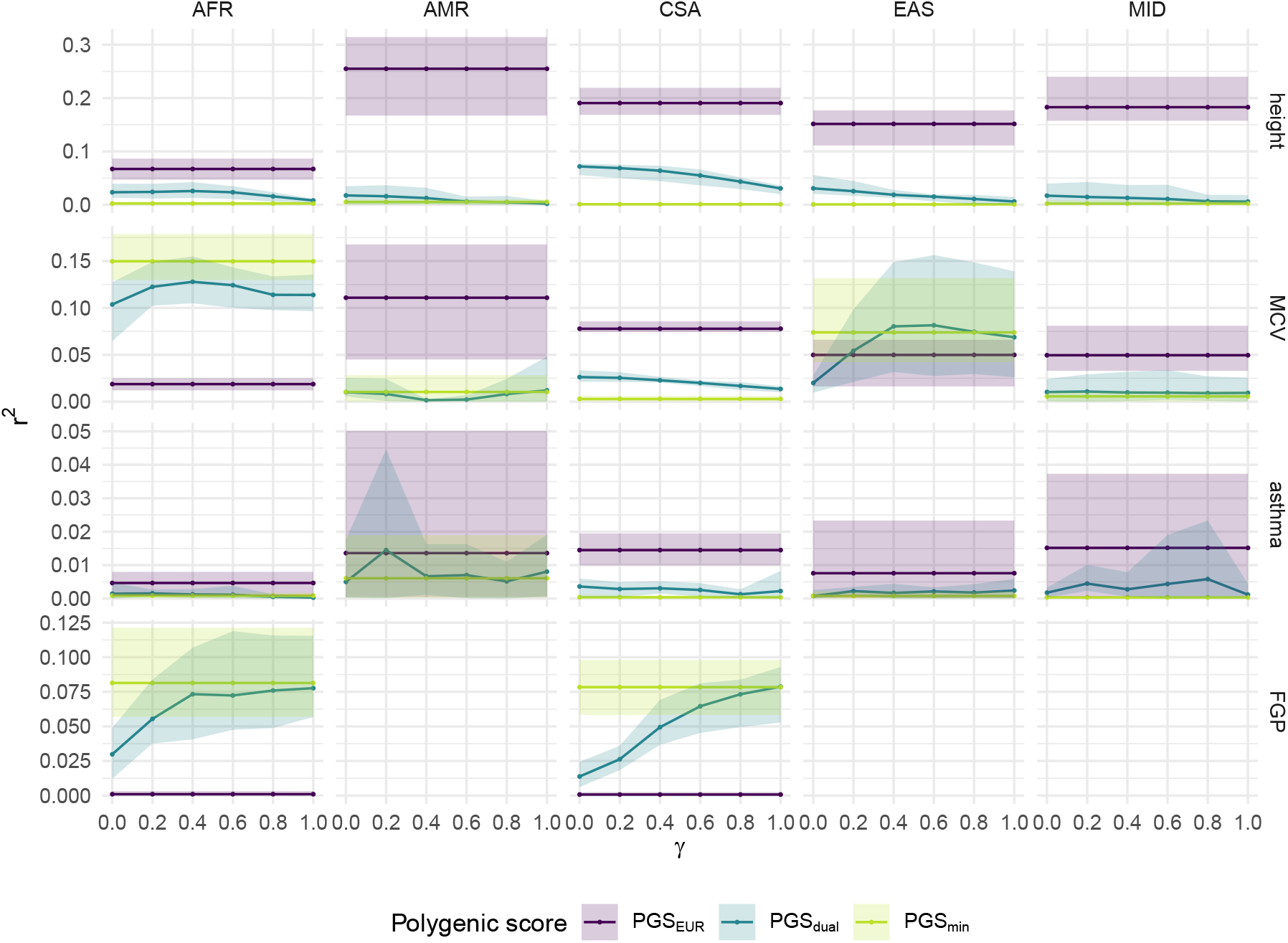
Partial *r*^2^ for PGS_EUR_, PGS_dual_, and PGS_min_ on four traits in UK Biobank for five minority-ancestry groups. The single-ancestry scores were estimated using a standard, unweighted LASSO. The dual-ancestry scores were constructed using an importance weighted LASSO with various degrees of reweighting *γ*. Error bars correspond to the range across five cross-validation rounds. The four traits considered are height, MCV, asthma, and FGP. We used inferred genetic ancestry labels from Pan-UKBB, with participants divided into six groups: European ancestry (EUR), African ancestry (AFR), Admixed American ancestry (AMR), Central/South Asian ancestry (CSA), East Asian ancestry (EAS), and Middle Eastern ancestry (MID). Analyses were not run on FGP for AMR, EAS and MID ancestry groups as the number of cases was fewer than 50 in each group.

### Differences in trait architecture explain variable performance by ancestry

Finally, to investigate the reasons why optimal training approaches vary across traits, we investigated the contribution of variants at different allele frequencies to variance explained. Specifically, we measured the partial *r*^2^ for different subsets of variants grouped by minor allele frequency (MAF) in a given ancestry group. We grouped variants into four sets: ‘rare’ (MAF ≤ 1%), ‘uncommon’ (1% < MAF ≤ 5%), ‘intermediate’ (5% < MAF ≤20%), and ‘common’ (MAF > 20%). Given a set of variants 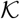 and the genotype matrix of the test set *X*, let 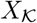 denote the submatrix given by the columns of *X* that are in 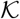. For a set of variants 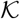, we define 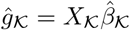 to be the score associated with the variant set 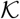, where 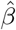 is the vector of variant effects obtained from the LASSO. We then defined the partial *r*^2^ attributable to variant set 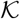 to be the difference in *r*^2^ between models

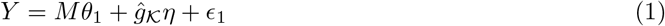

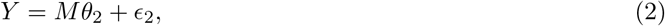

where *M* is the matrix of covariates (age, sex, the first ten genetic principal components (PCs), and interactions between sex and the ten genetic PCs) for the test set. We note that because we are considering the same set of variants for each trait, observed differences among traits in composition likely reflect differences in trait architecture, rather than differences in how the studies were performed.

Figure 6 illustrates the contribution to variance explained by different segments of the allele frequency spectrum for the PGS for mean corpuscular volume and height. We observe striking differences between these traits in the contribution of different allele frequency segments. For MCV in African-ancestry individuals, the majority of the variance explained in each of the polygenic scores was due to variants that were intermediate or common in African-ancestry individuals (see top row, LHS). Notably, over half the contribution to variance explained of the African-ancestry and PGS_dual_ could be attributed to variants that have MAF < 5% in the European-ancestry subgroup (second row, LHS). Conversely, variance explained in the European-ancestry subgroup was driven by variants that are intermediate or common in European-ancestry individuals (fourth row, LHS). And approximately one quarter of the variance explained could be attributed to variants with MAF < 5% in African-ancestry individuals.

**Figure 6:**
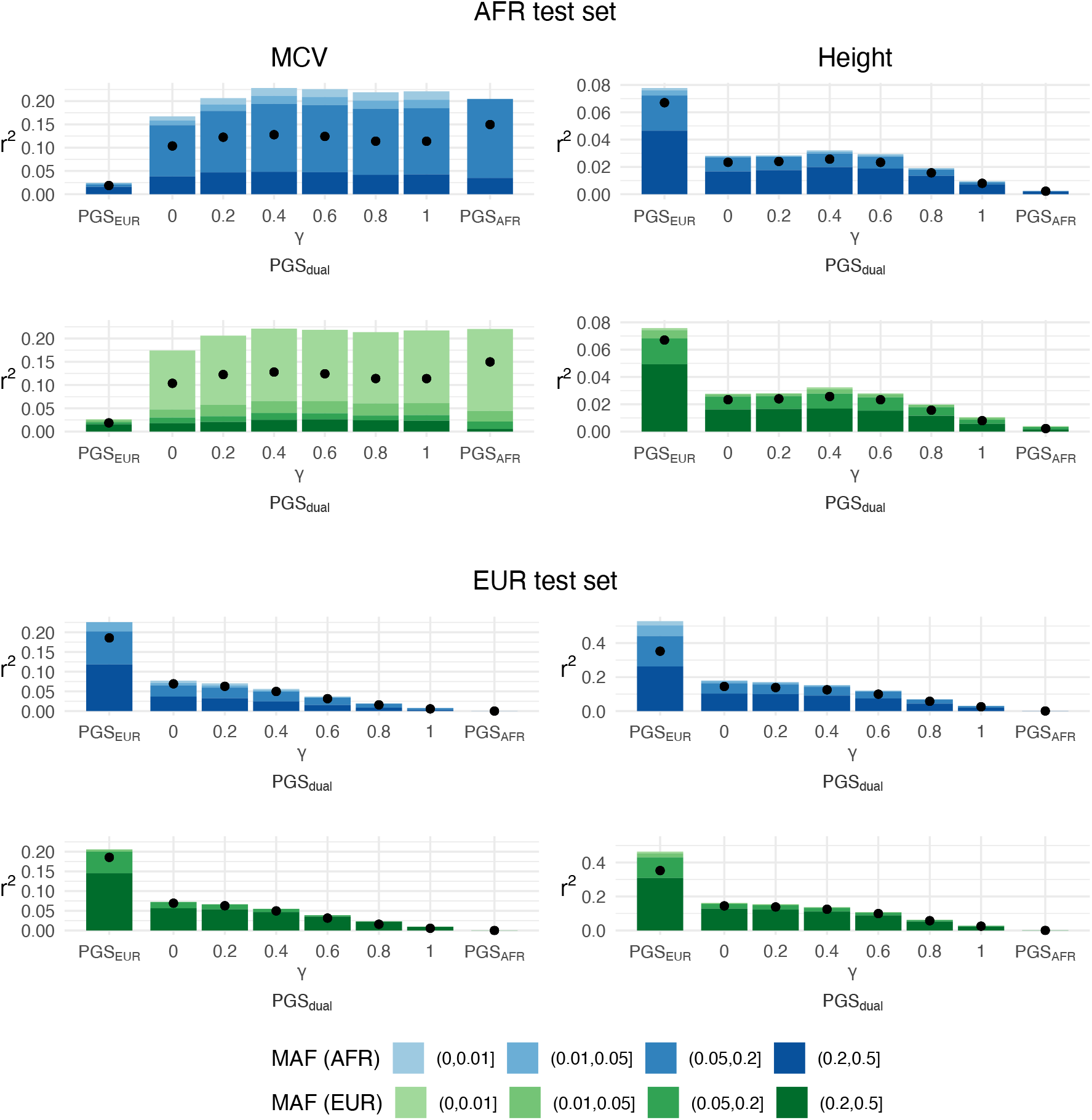
Allele frequency composition of variance explained by PGS for mean corpuscular volume (left) and height (right) in a African-ancestry test set (top) and a European-ancestry test set (bottom). The black dots represent partial *r*^2^ for all the variants, i.e. the entire polygenic score. Variants were grouped according to their minor allele frequency in African-ancestry individuals (blue palette) or in European-ancestry individuals (green palette). Each bar represents the sum of the partial *r*^2^ values for each subset of variants in a given polygenic score. Note that the height of the bar is generally higher than corresponding dot due to linkage disequilibrium between variants.

In contrast, for height we find that variance explained in European-ancestry individuals has an allele frequency decomposition with a slightly greater contribution of variants that are common in European-ancestry individuals (bottom row, RHS). But, variance explained in African-ancestry individuals is dominated by variants that are common (MAF > 5%) in both populations (top two rows, RHS). The effect of reweighting follows a similar pattern; for MCV shifting variance explained towards variants that are higher-frequency in the African-ancestry individuals (and considerably rarer in European-ancestry individuals) and, for height, tending to favour variants that are common in both groups.

These results suggest differences in genetic architecture between the two traits. For mean corpuscular volume, the variance explained by PGS_dual_ and PGS_AFR_ could largely be attributed to variants that were relatively common in African-ancestry individuals but rare in European-ancestry individuals. Since the number of African-ancestry individuals in these training sets was relatively small, this suggests that the corresponding effect sizes are comparatively large. On the other hand, the variance explained by each of the height PGS could be attributed to variants that were common in both European-ancestry and African-ancestry individuals. We found similar patterns for asthma and FGP, and for the other minority ancestries (SI Figures 2-10). Specifically, for traits and ancestries where PGS_min_ outperformed the PGS_EUR_, more variance could be attributed to variants that are more common in the minority ancestry group. We also found substantial differences across traits in the implied liability variance between PGS_EUR_ and PGS_min_ (Supplementary Table 1). Such differences in trait architecture are indicative of variation in the evolutionary and selective forces that have shaped trait variation [34, 35, 36].

## Discussion

The lack of diversity in human genetic studies has been brought into focus by a number of articles (e.g. [37, 38, 9, 39]), revealing that around 86% of all GWAS participants are of European descent. In fact, despite calls for more diversity in genomic studies, this Eurocentric bias appears to be getting worse: in 2019 the proportion of samples in the GWAS Catalog [40] from European-ancestry individuals was 78% [39]. In the context of polygenic scores, this bias has been reflected in the lack of transferability of scores across ancestries: PGS developed using European-ancestry samples tend to perform poorly in non-European ancestry test sets [7, 9, 8]. Correspondingly, our results found that, for a range of traits, PGS estimated using European-ancestry individuals performed relatively poorly on other non-European ancestry groups.

Recent improvements in predictive performance across a number of traits indicate the potential clinical utility of PGS [5, 6, 3, 2]. By more accurately quantifying an individual’s risk for a certain disease or trait, PGS may enhance the capability to predict disease development, progression, and recurrence, and to deploy more targeted preventive and therapeutic interventions [2, 41]. The disparity in PGS accuracy across ancestry groups, however, raises serious technical, clinical and ethical issues, with likely substantial impacts on health inequalities if left unaddressed [9].

Recent investigations into statistical methods have sought to construct improved PGS using the available data with the aim of reducing this disparity in predictive accuracy. Previous studies on the lack of transferability of PGS have generally estimated scores using summary statistics from genome-wide association studies of single-ancestry populations [7, 9, 8]. These summary statistic approaches are often highly efficient computationally and typically achieve highly competitive predictive performance relative to full genotype approaches [42, 43, 44, 32]. A number of methods to construct PGS by combining GWAS results from distinct ancestry groups have recently been developed, demonstrating improved prediction accuracy on the minority-ancestry group [20, 22, 21].

However, the question of how to optimally construct PGS directly from multiple-ancestry training data has largely been unexplored. That is, to maximise predictive accuracy for a minority-ancestry group, is it better to employ a PGS trained on just the minority-ancestry individuals or a combination of the minority-ancestry individuals and the majority-ancestry individuals? To investigate this, we compared the prediction accuracy of PGS trained on different subsets of the available data. This was possible due to the availability of individual-level genetic data in UK Biobank, accompanied by a wide range of phenotypic measurements. This in turn allowed us to vary the number of individuals of each ancestry in each training set and thus to assess the effect of sample size and composition on PGS predictive predictive performance for a range of quantitative and binary traits.

Our primary finding is that, in terms of minority-ancestry prediction performance, traits vary substantially in their optimal strategy for combining data across ancestries. Through simulation, we have shown that a PGS trained on a small minority-ancestry training set may outperform a much larger European-ancestry training set. Moreover, there are plausible regions of parameter space, notably where effect sizes are correlated across ancestry groups but not identical, where reweighting strategies can boost prediction performance using a multiple-ancestry training set. However, when applied to empirical data from the UK Biobank, relatively weak benefit from reweighting was observed. Rather, traits fell into two broad categories for which the minority-ancestry PGS outperformed the majority-ancestry PGS, or vice versa. For those in the first category, the most valuable training data was the small dataset from the minority group alone (and where performance decreases by the inclusion of any individuals of the majority - here European-ancestry). For those in the second category (such as height), the best performance in the underrepresented group was achieved by training in the much larger majority group, though note this still represented a substantial decrease in absolute performance relative to the majority European-ancestry group. When the minority group was African-ancestry individuals, the first category, though smaller, included FGP and important biomarkers such as MCV. Moreover, for a given trait, the optimal strategy varied by ancestry. For MCV for example, PGS_EUR_ outperformed the PGS_min_ for Admixed American-, Central/South Asian- and Middle Eastern-ancestry groups, while the reverse was true for the African-ancestry group. For the East Asian-ancestry group, the PGS displayed similar performance.

These findings raise the question of *why* the optimal strategy varies across traits and ancestry groups. Recent studies have ascribed the overall lack of PGS transferability to a number of factors, including population-level differences in allele frequencies, LD patterns, location of causal variants, and effect sizes [13, 14, 10, 9, 11, 12]. We found that the variability across traits could be at least partially explained by differences in the allele frequency pattern of causal variants or, more precisely, variants in LD with a causal variant. Specifically, we found that for some traits, such as height, most predictive utility could be attributed to variants that were relatively common (MAF > 5%) in both European- and African-ancestry individuals. For other traits, such as MCV, the predictive utility could be attributed to variants that were common in African-ancestry individuals but rare in European-ancestry individuals. This points to why PGS_EUR_ for MCV performed much worse than the corresponding PGS_AFR_, despite being trained on a much larger number of individuals.

Our findings highlight the limitations of statistical corrections alone in reducing the disparities of PGS performance between ancestries. There are numerous sources of potential bias that adversely affect the application of an analytical pipeline to individuals of minority and/or underrepresented groups, including problem selection, data collection, outcome definition, model development and lack of real-world impact assessments and considerations [45]. Therefore, statistical solutions must be considered in combination with efforts to address the global health research funding gap [46], diversity of the bioinformatics workforce [47] and assessing the impact and translation of analyses to real-world data [48] in addition to efforts to increase the number of non-European-ancestry participants in genetic research. Each of these partial solutions, in combination, provide essential contributions to reduce and remove opportunities for negative effects on health inequalities, particular amongst those from different ancestry groups [49].

It is becoming increasingly clear that the vast disparities in PGS performance can only be bridged by improving the diversity of human genetic datasets [9]. An important consideration surrounds future sampling strategy: whose genomes should we aim to sequence to reduce existing disparities in polygenic prediction accuracy? Our findings indicate that increasing the number of samples from minority-ancestry groups can lead to significant improvements in prediction performance. However, we note that targeting data collection from minority-ancestry groups is logistically difficult given that inferring genetic ancestry requires at least some genomic data to begin with, for example from an ancestry-informative marker set [50]. Moreover, even within the same ancestry group, PGSs do not always generalise across other characteristics such as age, sex and socio-economic status [51], pointing to the need of more diversity across multiple axes both alongside and within genetic ancestry.

Perhaps counter-intuitively, with more diverse data, statistical tools such as importance reweighting may eventually play a more important role as we seek to boost predictive utility by using all the available data. Reweighting strategies have the benefit of overcoming the lack of universal definitions of race, ethnicity and ancestry, which causes considerable confusion and imprecision [52, 53]. The categorisation of individuals into discrete ancestry groups to explain differences between behaviours and exposures may be unhelpful [54], whereas continuous representations of genetic ancestry [55, 56] enable identification of areas of genetic ancestry where performance is stronger or weaker as well as how the trait itself varies across ancestry. Ultimately, approaches to genetic prediction must acknowledge both the many similarities of human biology, but also the differences in history, cultural heritage, exposure, and behaviour that can lead to certain factors being of greater relevance for particular groups of individuals.

## Materials and Methods

### PGS Estimation using the LASSO

To construct PGSs from full genotype data, we use L1-penalised regression, also known as the LASSO [28]. The LASSO has previously been used in the context of genetic prediction (see for example [29, 30, 31, 32, 12] and references therein) and is suitable for high-dimensional problems where one expects the number of non-zero predictors to be small relative to the number of total predictors. Although there exist other full genotype PGS methods such as linear mixed models (see e.g. [57]), we focus on the LASSO largely for its computational efficiency. We provide an analysis of predictive performance versus sample size, focusing on differences in predictive performance between European-ancestry individuals and non-European-ancestry individuals.

We first briefly recap the LASSO algorithm for constructing polygenic scores. Let *n* be the number of individuals in the training set. Denote 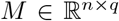 to be the matrix of *q* covariates, *X* to be the *n* × *p* genotype matrix, and *y* to be the *n*-vector of observed phenotype values. We assume a linear relationship between the phenotype and predictors,

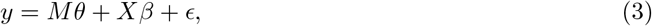

where *θ* is a *q*-vector of covariate effects, *β* is a *p*-vector of variant effects, and *ϵ* is an environmental noise term with mean 0.

The LASSO aims to minimise the objective function,

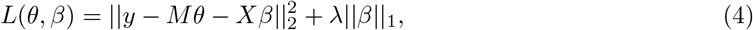

with respect to (*θ, β*), where *λ* is a regularisation parameter. The purpose of the ∥*β*∥_1_ term is to penalise large values of |*β*| and thus encourage *sparse* solutions, that is, solutions with a relatively low number of non-zero coefficients. Note that the parameters *θ* are not penalised. The choice of *λ* controls the degree of penalisation with higher values of *λ* encouraging smaller values of *β*. To select *λ*, we applied an 80-20 training-validation split to the training set, generating a path of solutions for a range of values of *λ* using the training split, and then selecting the value of *λ* that maximises *r*^2^ (for quantitative traits) or the Area Under the Curve (AUC; for binary traits) on the validation split.

We optimise (4) using the R packages `glmnet` [58] and `snpnet` [32] on simulated and real data respectively. The `snpnet` package is an extension of `glmnet` designed to interface directly with the PLINK software [59, 60] to handle large-scale single nucleotide polymorphism datasets such as the genotype data in UK Biobank. Using the default package settings and following the guidance in [32], we did not standardise the genotypes before fitting.

#### Multiple-ancestry Datasets

The above description assumes that the genetic effects *β* are the same for all individuals. While this assumption may be reasonable within a single ancestry group, it is unlikely to hold more generally. We consider a hypothetical model that allows genetic effects to vary among individuals. To this end, we introduce a low-dimensional latent variable *z* representing the ancestry of a given individual. For an individual *i*,

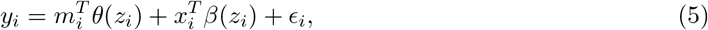

where (*θ*(·), *β*(·)) are now *functions* parameterised by *z*. The only assumption we make about these functions is one of *continuity*, so that individuals with similar ancestry have similar covariate and genetic effects. Otherwise, we do not make any explicit statements regarding the form of the functions.

In this paper, we consider the special case with *J* = 2 ancestry groups, such that *z_i_* ∈ {1, 2}. We denote (*θ*(*z_i_*), *β*(*z_i_*)) = (*θ*^(*j*)^, *β*^(*j*)^) for an individual *i* in ancestry group *j*. Using superscripts to denote the ancestry group (for example, *y*^(*j*)^ is the vector of observed phenotypes for group *j*),

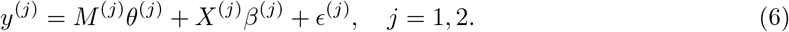

Denoting *n_j_* to be the number of individuals of ancestry *j* in the training set, we assume *n*_1_ > *n*_2_ to reflect a European-ancestry dominated training set. Applying the standard LASSO to such a training set, we would expect the estimate 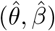 to be closer to (*θ*^(1)^, *β*^(1)^) than to (*θ*^(2)^, *β*^(2)^). All else being equal, this would result in better predictive performance for test individuals in group 1.

#### Importance Reweighting

We return for a moment to the more general model in Equation (5). Suppose we wish to construct a polygenic score for a test set with ancestry distribution *π^TE^* (*z*) given a training set of observations 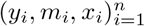 from individuals with ancestry *z*_1_,…, *z_n_*, how should one obtain a single estimate for (*θ, β*)? Our proposed approach is to use *importance reweighting*, a statistical technique commonly used in survey sampling [24]. Importance reweighting aims to account for distributional differences between the sampling population and the population of interest by assigning weights to each of the samples in the training data. Here, we assign each of the training individuals a weight *w_i_* that quantifies the ‘similarity’ of the ancestry variable *z_i_* to the test set ancestries, with individuals who have a similar ancestry to the test set assigned a higher weight. More concretely, assuming an ancestry distribution *π^TR^*(·) on the training set, we set

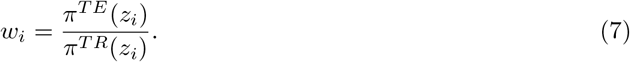

Note that we do not have access to the distributions *π^TR^*, *π^TE^* so in practice we must approximate (7). Given weights w_i_, to estimate (*θ, β*), we minimise the following *weighted* LASSO objective function:

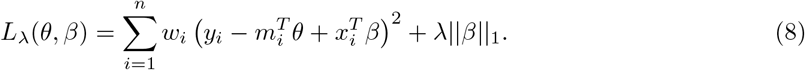

For the special case with two ancestry groups (Eqn. 6), we consider importance weights *w*_1_, *w*_2_ such that *w*_2_ ≥ *w*_1_ with the aim of improving estimates for group 2, i.e. the underrepresented group. Specifically, we use weights of the form:

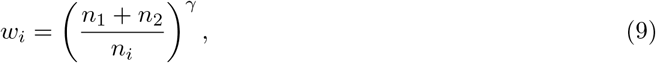

where *γ* ∈ [0, 1] is a hyperparameter that controls the degree of reweighting. We normalised the weights so that *n*_1_*w*_1_ + *n*_2_*w*_2_ = *n*_1_ + *n*_2_. With these weights, we thus minimise the objective function,

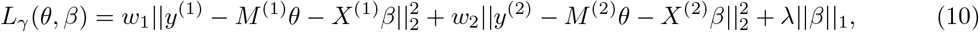

with respect to (*θ, β*).

### Simulation Study

To evaluate the effect of importance reweighting on PGS performance under various settings, we undertook a simulation study. We used the simulation engine `msprime` [25] in the standard library of population genetic simulation models `stdpopsim` v0.1.2 [26] to generate African-ancestry and European-ancestry genotypes, following a similar simulation framework to that used by Martin et al. [7]. For each ancestry group, we generated a total of 200 000 genotypes for chromosome 20, based on a three-population ‘out-of-Africa’ demographic model [61], using a mean mutation rate of 1.29 × 10^-8^ and a recombination map of GRCh37. This recombination map is from the Phase II Hapmap project and is based on 3.1 million genotyped SNPs from 270 individuals across four populations (Nigeria, Beijing, Japan, northern and western Europe) [62]. Note that this particular simulation model also generates genotypes for East Asian individuals but we do not use these in our analysis. To reduce the computational burden, we used only the first 10% of the chromosome, applying a minor allele frequency (MAF) threshold to filter out any SNPs that had a MAF of less than 1% in each ancestry group, resulting in a total of 189, 765 variants.

We simulated phenotypes from these genotypes assuming a normal linear model,

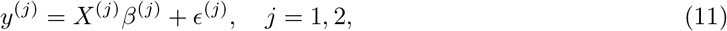

where 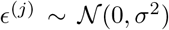. The noise variance parameter *σ*^2^ controls the SNP heritability 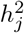 for each ancestry group and was chosen to yield an average SNP heritability of 0.3 across the groups. To reflect the sparsity of genetic effects, we randomly selected *p*_0_ = 100 of the *p* = 189, 765 variants to be causal. Motivated by Trochet et al. [63], we drew the causal effect sizes from a bivariate normal distribution to model the similarity of genetic effects between ancestries. Denoting 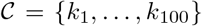 to be the indices of the causal SNPs, and 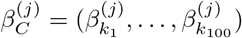 we have the equation

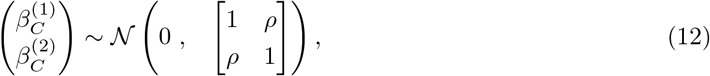

where *ρ* is the correlation between the ancestry-specific genetic effects. We set 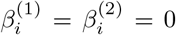 for *i* ∉ *C*. For each value of *ρ* = 0.5, 0.6,…, 0.9, we repeated the above procedure 10 times to generate 10 quantitative traits.

To investigate the effect of sample size, for each trait we created five training sets with *n_EUR_* = 18 000 randomly selected individuals of European ancestry and *n_AFR_* = 2 000, 4 500, 7 714, 12 000, 18 000 randomly selected individuals of African ancestry. For each of these training sets, we computed weights 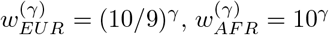 and where *γ* is a hyperparameter controlling the degree of reweighting. Note that *γ* = 1 corresponds to inverse proportion reweighting when *n_AFR_* = 2 000 and *n_EUR_* = 18 000. We normalised the weights to ensure that their sum equalled *n*, in line with the unweighted case.

To assess predictive performance for each ancestry, we constructed an African-ancestry test set and a European-ancestry test set by randomly selecting 2 000 individuals from each ancestry out of those not included in the training set. We used the proportion of variance explained, denoted *r*^2^, as the measure of predictive performance,

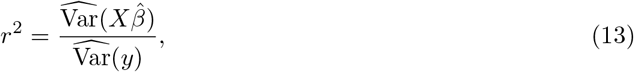

where 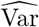 denotes the sample variance. Note that this is equivalent to the partial *r*^2^ relative to an intercept-only covariate model. For *ρ* = 0.5, 0.6,…, 0.9, we repeated the above process 20 times with different randomly selected individuals in the training and test sets to calculate a mean *r*^2^ for each trait.

### UK Biobank

The UK Biobank is a large-scale cohort study with genomic and phenotypic data collected on approximately 500 000 individuals aged 40-69 at the time of recruitment [23]. We used inferred genetic ancestry labels made available by the Pan-UKBB initiative [27]. These labels were derived based on reference data from the 1000 Genomes Project [64] and the Human Genome Diversity Project (HGDP) [65]. Briefly, principal component analysis (PCA) was performed on unrelated individuals in the combined reference dataset, and a random forest classifier was trained on the first 6 principal components (PCs) using continental ancestry metadata. UK Biobank individuals were then projected onto this reference PC space and given initial ancestry assignments based on the random forest classifier. These assignments were then adjusted by removing outliers. Full details can be found on the Pan-UKBB website [27].

#### Phenotypes

We investigated a range of quantitative and binary traits available in our UK Biobank project (UKB Application Number 12788). To select these, we applied the following procedure. We first filtered out traits with estimated SNP heritability of less than 5% or a ‘low’ confidence rating for the SNP heritability estimate, as reported on the Neale Lab UKB SNP-Heritability Browser [66, 67]. To ensure that the traits under investigation were not highly correlated with each other, we used genetic correlation estimates available on the Neale Lab UKBB Genetic Correlation Browser [68, 69]. Specifically, working from the most heritable traits to the least heritable, we iteratively removed a trait if it had an estimated genetic correlation of more than 0.5 with any remaining trait of higher heritability. Of the remaining traits, we selected the top 10 most heritable quantitative traits: height, mean corpuscular volume (MCV), body mass index (BMI), eosinophill percentage, erythrocyte distribution width (erythrocyte), lymphocyte count, monocyte count, mean platelet volume (MPV), platelet crit, and high light scatter reticulocyte count (reticulocyte). We also selected the top 5 most heritable binary traits: asthma, female genital prolapse (FGP), atrial fibrillation (AFib), diverticular disease of the intestine (DDI), and hypothyroidism. For our African-ancestry analyses, we investigated all 15 of these traits, while for the other minority-ancestry analyses, we focused in on 4 traits: height, mean corpuscular volume, female genital prolapse, and asthma.

#### Genotypes and covariates

We used imputed genotypes from UK Biobank, filtering out individuals and variants according to quality-control metrics used by the Pan-UKBB initiative [27]. Firstly, we removed individuals who were identified as displaying sex chromosome aneuploidy. We also removed individuals who were flagged as related by Pan-UKBB. Within each ancestry group, related individuals were identified using PC-Relate [71] with *k* = 10 and a minimum individual MAF of 5%. We filtered out variants that were not deemed to be of ‘high quality’ according to Pan-UKBB, retaining those with an INFO score of at least 0.8 and with an allele count of at least 20 per population. We also removed variants that had a MAF of less than 1% in both the European-ancestry group and the minority-ancestry group under analysis. To control for population structure, we included the following covariates in our model: age, sex, the first ten genetic principal components (PCs), and interactions between sex and the ten genetic PCs.

#### Construction of Training Sets

To assess the effect of sample size and composition on PGS performance, we used subsets of the above data as training sets, controlling the number of European-ancestry and minority-ancestry individuals. The remaining individuals were then used as a held-out test set. For each trait, we constructed three types of training sets using individuals with non-missing data for that particular trait. The first two types of training sets were single-ancestry datasets, one consisting solely of European-ancestry individuals and the other consisting solely of minority-ancestry individuals.

1. *European-ancestry* A random sample comprising 80% of the quality-controlled European-ancestry individuals.
2. *Minority-ancestry* A random sample comprising 80% of the respective quality-controlled minority-ancestry individuals.
3. *Dual-ancestry* This set consisted of both minority-ancestry individuals and European-ancestry individuals. The basic form of this training set was made up of the minority-ancestry training set described above, combined with European-ancestry individuals so that the proportion of minority-ancestry individuals was 10%. The European-ancestry individuals were matched to the minority-ancestry individuals on age and sex. We also considered variations of this training set by removing a proportion of either the European-ancestry individuals or the minority-ancestry individuals (see Results for more details).

For each dataset, we further removed variants with a minor allele frequency of less than 0.1% or a missing genotype call rate of less than 5%. Note that, as a result, the sets of variants generally differed by training set. For the single-ancestry datasets, we used the standard, unweighted LASSO to construct the PGS. For the dual-ancestry datasets, we used the weighted LASSO with *γ* = 0, 0.2, 0.4, 0.6, 0.8, 1. Note that *γ* = 0 corresponds to the unweighted LASSO.

#### Predictive Accuracy

We evaluated predictive accuracy of each PGS by ancestry group using individuals that were not included in the corresponding training sets. To assess the genetic predictive accuracy of a PGS, we calculated the partial *r*^2^ attributable to the PGS, relative to a covariate-only model, following Martin et al. [9]. Specifically, we fit the following nested linear models (or the equivalent logistic models for binary traits),

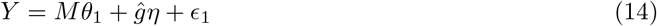

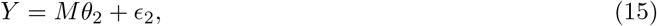

where *M* is the covariate matrix and g is the vector consisting of polygenic scores for each individual in the test set. We took the partial *r*^2^ (or the Cox and Snell pseudo-*r*^2^ for binary traits) between models 14 and 15 as our measure of predictive accuracy. To obtain more reliable estimates, we performed 5-fold cross-validation. That is, we repeated five times the process of training set construction and PGS estimation to obtain five estimates of partial *r*^2^, and report the mean and the range of these estimates.

## Data Availability

This research has been conducted using data from UK Biobank, a major biomedical database, under application 12788 [23]. We used genetic correlation estimates from the Neale Lab UKBB Genetic Correlation Browser [68, 69] and heritability estimates from the Neale Lab UKB SNP-Heritability Browser [66, 67]. We use ancestry-specific heritability estimates and inferred genetic ancestry labels for UK Biobank individuals made available by the Pan-UKBB initiative [27].

## Acknowledgements

BCLL, GM and CCH were supported by the UK Engineering and Physical Sciences Research Council through the Bayes4Health programme [Grant number EP/R018561/1]. BCLL also gratefully acknowledges funding from Jesus College, Oxford. CCH was also supported by The Alan Turing Institute, Health Data Research UK, the Medical Research Council UK, and AI for Science and Government UK Research and Innovation (UKRI). GM was supported by the Wellcome Trust [Grant Number 100956/Z/13/Z] and the Li Ka Shing Foundation. The computational aspects of this research were supported by the Wellcome Trust Core Award Grant Number 203141/Z/16/Z and the NIHR Oxford BRC. The views expressed are those of the authors and not necessarily those of the NHS, the NIHR or the Department of Health.

## Notes

### Competing Interest Statement

G.M. is a director of and shareholder in Genomics PLC, and is a partner in Peptide Groove LLP.

### Summary of Updates

Substantial revisions, including analysis of more traits and more ancestries, use of imputed data instead of genotyped data, larger simulation study.

## References

[1] Nilanjan Chatterjee, Jianxin Shi, and Montserrat García-Closas. “Developing and evaluating polygenic risk prediction models for stratified disease prevention”. In: Nature Reviews Genetics 17.7 (July 2016), pp. 392–406. ISSN: 1471-0064. DOI: 10.1038/nrg.2016.27. URL: https://doi.org/10.1038/nrg.2016.27.

[2] Ali Torkamani, Nathan E. Wineinger, and Eric J. Topol. “The personal and clinical utility of polygenic risk scores”. In: Nature Reviews Genetics 19.9 (Sept. 2018), pp. 581–590. ISSN: 1471-0064. DOI: 10.1038/s41576-018-0018-x. URL: https://doi.org/10.1038/s41576-018-0018-x.

[3] Amit V. Khera et al. “Genome-wide polygenic scores for common diseases identify individuals with risk equivalent to monogenic mutations”. In: Nature Genetics 50.9 (Sept. 2018), pp. 1219–1224. ISSN: 1546-1718. DOI: 10.1038/s41588-018-0183-z. URL: https://doi.org/10.1038/s41588-018-0183-z.

[4] Joshua W. Knowles and Euan A. Ashley. “Cardiovascular disease: The rise of the genetic risk score”. In: PLOS Medicine 15.3 (Mar. 2018), pp. 1–7. DOI: 10.1371/journal.pmed.1002546. URL: https://doi.org/10.1371/journal.pmed.1002546.

[5] Paige Maas et al. “Breast Cancer Risk From Modifiable and Nonmodifiable Risk Factors Among White Women in the United States”. In: JAMA Oncology 2.10 (Oct. 2016). ISSN: 2374-2437. DOI: 10.1001/jamaoncol.2016.1025. URL: https://doi.org/10.1001/jamaoncol.2016.1025.

[6] Seth A. Sharp et al. “Development and Standardization of an Improved Type 1 Diabetes Genetic Risk Score for Use in Newborn Screening and Incident Diagnosis”. In: Diabetes Care 42.2 (Jan. 2019), pp. 200–207. ISSN: 0149-5992. DOI: 10.2337/dc18-1785. URL: https://doi.org/10.2337/dc18-1785.

[7] Alicia R. Martin et al. “Human Demographic History Impacts Genetic Risk Prediction across Diverse Populations”. In: The American Journal of Human Genetics 100.4 (2017), pp. 635–649. ISSN: 0002-9297. DOI: https://doi.org/10.1016/j.ajhg.2017.03.004. URL: https://www.sciencedirect.com/science/article/pii/S0002929717301076.

[8] L. Duncan et al. “Analysis of polygenic risk score usage and performance in diverse human populations”. In: Nature Communications 10.1 (July 2019), p. 3328. ISSN: 2041-1723. DOI: 10.1038/s41467-019-11112-0. URL: https://doi.org/10.1038/s41467-019-11112-0.

[9] Alicia R. Martin et al. “Clinical use of current polygenic risk scores may exacerbate health disparities”. In: Nature Genetics 51.4 (Apr. 2019), pp. 584–591. ISSN: 1546-1718. DOI: 10.1038/s41588-019-0379-x. URL: https://doi.org/10.1038/s41588-019-0379-x.

[10] Marco Scutari, Ian Mackay, and David Balding. “Using Genetic Distance to Infer the Accuracy of Genomic Prediction”. In: PLOS Genetics 12.9 (Sept. 2016), pp. 1–19. DOI: 10.1371/journal.pgen.1006288. URL: https://doi.org/10.1371/journal.pgen.1006288.

[11] Ying Wang et al. “Theoretical and empirical quantification of the accuracy of polygenic scores in ancestry divergent populations”. In: Nature Communications 11.1 (July 2020), p. 3865. ISSN: 2041-1723. DOI: 10.1038/s41467-020-17719-y. URL: https://doi.org/10.1038/s41467-020-17719-y.

[12] Florian Privé et al. “Portability of 245 polygenic scores when derived from the UK Biobank and applied to 9 ancestry groups from the same cohort”. In: The American Journal of Human Genetics 109.1 (2022), pp. 12–23. ISSN: 0002-9297. DOI: https://doi.org/10.1016/j.ajhg.2021.11.008. URL: https://www.sciencedirect.com/science/article/pii/S0002929721004201.

[13] Christopher S. Carlson et al. “Generalization and Dilution of Association Results from European GWAS in Populations of Non-European Ancestry: The PAGE Study”. In: PLOS Biology 11.9 (Sept. 2013), pp. 1–11. DOI: 10.1371/journal.pbio.1001661. URL: https://doi.org/10.1371/journal.pbio.1001661.

[14] Brielin C. Brown et al. “Transethnic Genetic-Correlation Estimates from Summary Statistics”. In: The American Journal of Human Genetics 99.1 (July 2016). Publisher: Elsevier, pp. 76–88. ISSN: 0002-9297. DOI: 10.1016/j.ajhg.2016.05.001. URL: https://doi.org/10.1016/j.ajhg.2016.05.001.

[15] Paul W. Franks, Ewan Pearson, and Jose C. Florez. “Gene-Environment and Gene-Treatment Interactions in Type 2 Diabetes: Progress, pitfalls, and prospects”. In: Diabetes Care 36.5 (Apr. 2013). _eprint: https://diabetesjournals.org/care/article-pdf/36/5/1413/617153/1413.pdf, pp. 1413–1421. ISSN: 0149-5992. DOI: 10.2337/dc12-2211. URL: https://doi.org/10.2337/dc12-2211.

[16] Amy R. Bentley et al. “Multi-ancestry genome-wide gene–smoking interaction study of 387,272 individuals identifies new loci associated with serum lipids”. In: Nature Genetics 51.4 (Apr. 2019), pp. 636–648. ISSN: 1546-1718. DOI: 10.1038/s41588-019-0378-y. URL: https://doi.org/10.1038/s41588-019-0378-y.

[17] H3 Africa Consortium et al. “Enabling the genomic revolution in Africa”. In: Science 344.6190 (2014), pp. 1346–1348. DOI: 10.1126/science.1251546. eprint: https://www.science.org/doi/pdf/10.1126/science.1251546. URL: https://www.science.org/doi/abs/10.1126/science.1251546.

[18] Kelsey E. Grinde et al. “Generalizing polygenic risk scores from Europeans to Hispanics/Latinos”. In: Genetic Epidemiology 43.1 (2019), pp. 50–62. DOI: https://doi.org/10.1002/gepi.22166. eprint: https://onlinelibrary.wiley.com/doi/pdf/10.1002/gepi.22166. URL: https://onlinelibrary.wiley.com/doi/abs/10.1002/gepi.22166.

[19] Carla Marquez-Luna et al. “Multiethnic polygenic risk scores improve risk prediction in diverse populations”. In: Genetic Epidemiology 41.8 (2017), pp. 811–823. DOI: https://doi.org/10.1002/gepi.22083. eprint: https://onlinelibrary.wiley.com/doi/pdf/10.1002/gepi.22083. URL: https://onlinelibrary.wiley.com/doi/abs/10.1002/gepi.22083.

[20] Omer Weissbrod et al. “Leveraging fine-mapping and non-European training data to improve cross-population polygenic risk scores”. In: medRxiv (2021). DOI: 10.1101/2021.01.19.21249483. URL: https://www.medrxiv.org/content/early/2021/08/20/2021.01.19.21249483.

[21] Yunfeng Ruan et al. “Improving Polygenic Prediction in Ancestrally Diverse Populations”. In: medRxiv (2021). DOI: 10.1101/2020.12.27.20248738. URL: https://www.medrxiv.org/content/early/2021/08/24/2020.12.27.20248738.

[22] Taylor B. Cavazos and John S. Witte. “Inclusion of variants discovered from diverse populations improves polygenic risk score transferability”. In: Human Genetics and Genomics Advances 2.1 (2021), p. 100017. ISSN: 2666-2477. DOI: https://doi.org/10.1016/j.xhgg.2020.100017. URL: https://www.sciencedirect.com/science/article/pii/S2666247720300178.

[23] Clare Bycroft et al. “The UK Biobank resource with deep phenotyping and genomic data”. In: Nature 562.7726 (Oct. 2018), pp. 203–209. ISSN: 1476-4687. DOI: 10.1038/s41586-018-0579-z. URL: https://doi.org/10.1038/s41586-018-0579-z.

[24] Richard L Scheaffer et al. Elementary survey sampling. Cengage Learning, 2011.

[25] Jerome Kelleher, Alison M Etheridge, and Gilean McVean. “Efficient Coalescent Simulation and Genealogical Analysis for Large Sample Sizes”. In: PLoS Comput Biol 12.5 (May 2016), pp. 1–22. DOI: 10.1371/journal.pcbi.1004842. URL: http://dx.doi.org/10.1371%2Fjournal.pcbi.1004842.

[26] Jeffrey R Adrion et al. “A community-maintained standard library of population genetic models”. In: eLife 9 (June 2020). Ed. by Graham Coop et al., e54967. ISSN: 2050-084X. URL: https://doi.org/10.7554/eLife.54967.

[27] The Pan-UKBB team. 2020. URL: https://pan.ukbb.broadinstitute.org.

[28] Robert Tibshirani. “Regression Shrinkage and Selection Via the Lasso”. In: Journal of the Royal Statistical Society: Series B (Methodological) 58.1 (1996), pp. 267–288. DOI: https://doi.org/10.1111/j.2517-6161.1996.tb02080.x. URL:https://rss.onlinelibrary.wiley.com/doi/abs/10.1111/j.2517-6161.1996.tb02080.x.

[29] Patrik Waldmann et al. “AUTALASSO: an automatic adaptive LASSO for genome-wide prediction”. In: BMC Bioinformatics 20.1 (Apr. 2019), p. 167. ISSN: 1471-2105. DOI: 10.1186/s12859-019-2743-3. URL: https://doi.org/10.1186/s12859-019-2743-3.

[30] Sebastian Okser et al. “Regularized Machine Learning in the Genetic Prediction of Complex Traits”. In: PLOS Genetics 10.11 (Nov. 2014), pp. 1–9. DOI: 10.1371/journal.pgen.1004754. URL: https://doi.org/10.1371/journal.pgen.1004754.

[31] Florian Prive, Hugues Aschard, and Michael G. B. Blum. “Efficient Implementation of Penalized Regression for Genetic Risk Prediction”. In: Genetics 212.1 (2019), pp. 65–74. ISSN: 0016-6731. DOI: 10.1534/genetics.119.302019. eprint: https://www.genetics.org/content/212/1/65.full.pdf. URL: https://www.genetics.org/content/212/1/65.

[32] Junyang Qian et al. “A fast and scalable framework for large-scale and ultrahigh-dimensional sparse regression with application to the UK Biobank”. In: PLOS Genetics 16.10 (Oct. 2020), pp. 1–31. DOI: 10.1371/journal.pgen.1009141. URL: https://doi.org/10.1371/journal.pgen.1009141.

[33] Kevin J. Galinsky et al. “Estimating cross-population genetic correlations of causal effect sizes”. In: Genetic Epidemiology 43.2 (2019), pp. 180–188. DOI: https://doi.org/10.1002/gepi.22173. eprint: https://onlinelibrary.wiley.com/doi/pdf/10.1002/gepi.22173. URL: https://onlinelibrary.wiley.com/doi/abs/10.1002/gepi.22173.

[34] Yuval B. Simons et al. “A population genetic interpretation of GWAS findings for human quantitative traits”. In: PLOS Biology 16.3 (Mar. 2018), pp. 1–20. DOI: 10.1371/journal.pbio.2002985. URL: https://doi.org/10.1371/journal.pbio.2002985.

[35] Jian Zeng et al. “Signatures of negative selection in the genetic architecture of human complex traits”. In: Nature Genetics 50.5 (May 2018), pp. 746–753. ISSN: 1546-1718. DOI: 10.1038/s41588-018-0101-4. URL: https://doi.org/10.1038/s41588-018-0101-4.

[36] Armin P. Schoech et al. “Quantification of frequency-dependent genetic architectures in 25 UK Biobank traits reveals action of negative selection”. In: Nature Communications 10.1 (Feb. 2019), p. 790. ISSN: 2041-1723. DOI: 10.1038/s41467-019-08424-6. URL: https://doi.org/10.1038/s41467-019-08424-6.

[37] Anna C. Need and David B. Goldstein. “Next generation disparities in human genomics: concerns and remedies”. In: Trends in Genetics 25.11 (Nov. 2009), pp. 489–494. ISSN: 0168-9525. DOI: 10.1016/j.tig.2009.09.012. URL: https://doi.org/10.1016/j.tig.2009.09.012.

[38] Alice B. Popejoy and Stephanie M. Fullerton. “Genomics is failing on diversity”. In: Nature 538.7624 (Oct. 2016), pp. 161–164. ISSN: 1476-4687. DOI: 10.1038/538161a. URL: https://doi.org/10.1038/538161a.

[39] Segun Fatumo et al. “A roadmap to increase diversity in genomic studies”. In: Nature Medicine (Feb. 2022). ISSN: 1546-170X. DOI: 10.1038/s41591-021-01672-4. URL: https://doi.org/10.1038/s41591-021-01672-4.

[40] Annalisa Buniello et al. “The NHGRI-EBI GWAS Catalog of published genome-wide association studies, targeted arrays and summary statistics 2019”. In: Nucleic Acids Research 47.D1 (Nov. 2018), pp. D1005–D1012. ISSN: 0305-1048. DOI: 10.1093/nar/gky1120. eprint: https://academic.oup.com/nar/article-pdf/47/D1/D1005/27437312/gky1120.pdf. URL: https://doi.org/10.1093/nar/gky1120.

[41] Adebowale Adeyemo et al. “Responsible use of polygenic risk scores in the clinic: potential benefits, risks and gaps”. In: Nature Medicine 27.11 (Nov. 2021), pp. 1876–1884. ISSN: 1546-170X. DOI: 10.1038/s41591-021-01549-6. URL: https://doi.org/10.1038/s41591-021-01549-6.

[42] Luke R. Lloyd-Jones et al. “Improved polygenic prediction by Bayesian multiple regression on summary statistics”. In: Nature Communications 10.1 (Nov. 2019), p. 5086. ISSN: 2041-1723. DOI: 10.1038/s41467-019-12653-0. URL: https://doi.org/10.1038/s41467-019-12653-0.

[43] Timothy Shin Heng Mak et al. “Polygenic scores via penalized regression on summary statistics”. In: Genetic Epidemiology 41.6 (2017), pp. 469–480. DOI: https://doi.org/10.1002/gepi.22050. eprint: https://onlinelibrary.wiley.com/doi/pdf/10.1002/gepi.22050. URL: https://onlinelibrary.wiley.com/doi/abs/10.1002/gepi.22050.

[44] Bjarni J. Vilhjálmsson et al. “Modeling Linkage Disequilibrium Increases Accuracy of Polygenic Risk Scores”. In: The American Journal of Human Genetics 97.4 (Oct. 2015), pp. 576–592. ISSN: 0002-9297. DOI: 10.1016/j.ajhg.2015.09.001. URL: https://doi.org/10.1016/j.ajhg.2015.09.001.

[45] Irene Y. Chen et al. “Ethical Machine Learning in Healthcare”. In: Annual Review of Biomedical Data Science 4.1 (2021), pp. 123–144. DOI: 10.1146/annurev-biodatasci-092820-114757. eprint: https://doi.org/10.1146/annurev-biodatasci-092820-114757. URL: https://doi.org/10.1146/annurev-biodatasci-092820-114757.

[46] D. Vidyasagar. “Global notes: the 10/90 gap disparities in global health research”. In: Journal of Perinatology 26.1 (Jan. 2006), pp. 55–56. ISSN: 1476-5543. DOI: 10.1038/sj.jp.7211402. URL: https://doi.org/10.1038/sj.jp.7211402.

[47] Bas Hofstra et al. “The Diversity-Innovation Paradox in Science”. In: Proceedings of the National Academy of Sciences 117.17 (2020), pp. 9284–9291. ISSN: 0027-8424. DOI: 10.1073/pnas.1915378117. eprint: https://www.pnas.org/content/117/17/9284.full.pdf. URL: https://www.pnas.org/content/117/17/9284.

[48] Ziad Obermeyer et al. “Dissecting racial bias in an algorithm used to manage the health of populations”. In: Science 366.6464 (2019), pp. 447–453. DOI: 10.1126/science.aax2342. eprint: https://www.science.org/doi/pdf/10.1126/science.aax2342. URL: https://www.science.org/doi/abs/10.1126/science.aax2342.

[49] Roseann E. Peterson et al. “Genome-wide Association Studies in Ancestrally Diverse Populations: Opportunities, Methods, Pitfalls, and Recommendations”. In: Cell 179.3 (2019), pp. 589–603. ISSN: 0092-8674. DOI: https://doi.org/10.1016/j.cell.2019.08.051. URL: https://www.sciencedirect.com/science/article/pii/S0092867419310025.

[50] Kenneth K. Kidd et al. “Progress toward an efficient panel of SNPs for ancestry inference”. In: Forensic Science International: Genetics 10 (2014), pp. 23–32. ISSN: 1872-4973. DOI: https://doi.org/10.1016/j.fsigen.2014.01.002. URL: https://www.sciencedirect.com/science/article/pii/S1872497314000039.

[51] Hakhamanesh Mostafavi et al. “Variable prediction accuracy of polygenic scores within an ancestry group”. In: eLife 9 (Jan. 2020). Ed. by Ruth Loos, Michael B Eisen, and Paul O’Reilly, e48376. ISSN: 2050-084X. DOI: 10.7554/eLife.48376. URL: https://doi.org/10.7554/eLife.48376.

[52] Iain Mathieson and Aylwyn Scally. “What is ancestry?” In: PLOS Genetics 16.3 (Mar. 2020), pp. 1–6. DOI: 10.1371/journal.pgen.1008624. URL: https://doi.org/10.1371/journal.pgen.1008624.

[53] Tesfaye B. Mersha and Tilahun Abebe. “Self-reported race/ethnicity in the age of genomic research: its potential impact on understanding health disparities”. In: Human Genomics 9.1 (Jan. 2015), p. 1. ISSN: 1479-7364. DOI: 10.1186/s40246-014-0023-x. URL: https://doi.org/10.1186/s40246-014-0023-x.

[54] Morris W. Foster and Richard R. Sharp. “Race, Ethnicity, and Genomics: Social Classifications as Proxies of Biological Heterogeneity”. In: Genome Research 12.6 (2002), pp. 844–850. DOI: 10.1101/gr.99202. eprint: http://genome.cshlp.org/content/12/6/844.full.pdf+html. URL: http://genome.cshlp.org/content/12/6/844.abstract.

[55] Gillian M. Belbin et al. “Toward a fine-scale population health monitoring system”. In: Cell 184.8 (2021), 2068–2083.e11. ISSN: 0092-8674. DOI: https://doi.org/10.1016/j.cell.2021.03.034. URL: https://www.sciencedirect.com/science/article/pii/S0092867421003652.

[56] Anna C. F. Lewis et al. Getting Genetic Ancestry Right for Science and Society. 2021. arXiv: 2110.05987 [q-bio.PE].

[57] Xiang Zhou, Peter Carbonetto, and Matthew Stephens. “Polygenic Modeling with Bayesian Sparse Linear Mixed Models”. In: PLOS Genetics 9.2 (Feb. 2013), pp. 1–14. DOI: 10.1371/journal.pgen.1003264. URL: https://doi.org/10.1371/journal.pgen.1003264.

[58] Jerome H. Friedman, Trevor Hastie, and Rob Tibshirani. “Regularization Paths for Generalized Linear Models via Coordinate Descent”. In: Journal of Statistical Software 33.1 (2010), pp. 1–22. DOI: 10.18637/jss.v033.i01. URL: https://www.jstatsoft.org/index.php/jss/article/view/v033i01.

[59] Christopher C. Chang et al. “Second-generation PLINK: rising to the challenge of larger and richer datasets”. In: GigaScience 4.1 (Feb. 2015), p. 7. ISSN: 2047-217X. DOI: 10.1186/s13742-015-0047-8. URL: https://doi.org/10.1186/s13742-015-0047-8.

[60] Christopher C Chang and Shaun M Purcell. PLINK 1.9. www.cog-genomics.org/plink/1.9/.

[61] Ryan N. Gutenkunst et al. “Inferring the Joint Demographic History of Multiple Populations from Multidimensional SNP Frequency Data”. In: PLOS Genetics 5.10 (Oct. 2009), pp. 1–11. DOI: 10.1371/journal.pgen.1000695. URL: https://doi.org/10.1371/journal.pgen.1000695.

[62] The International HapMap Consortium. “A second generation human haplotype map of over 3.1 million SNPs”. In: Nature 449.7164 (Oct. 2007), pp. 851–861. ISSN: 1476-4687. DOI: 10.1038/nature06258. URL: https://doi.org/10.1038/nature06258.

[63] Holly Trochet et al. “Bayesian meta-analysis across genome-wide association studies of diverse phenotypes”. In: Genetic Epidemiology 43.5 (2019), pp. 532–547. DOI: https://doi.org/10.1002/gepi.22202. URL: https://onlinelibrary.wiley.com/doi/abs/10.1002/gepi.22202.

[64] Adam Auton et al. “A global reference for human genetic variation”. In: Nature 526.7571 (Oct. 2015), pp. 68–74. ISSN: 1476-4687. DOI: 10.1038/nature15393. URL: https://doi.org/10.1038/nature15393.

[65] Howard M. Cann et al. “A Human Genome Diversity Cell Line Panel”. In: Science 296.5566 (2002), pp. 261–262. DOI: 10.1126/science.296.5566.261b. URL: https://www.science.org/doi/abs/10.1126/science.296.5566.261b.

[66] Brendan K. Bulik-Sullivan et al. “LD Score regression distinguishes confounding from polygenicity in genome-wide association studies”. In: Nature Genetics 47.3 (Mar. 2015), pp. 291–295. ISSN: 1546-1718. DOI: 10.1038/ng.3211. URL: https://doi.org/10.1038/ng.3211.

[67] Neale Lab. 2019. URL: https://nealelab.github.io/UKBB_ldsc/h2_browser.html.

[68] Brendan Bulik-Sullivan et al. “An atlas of genetic correlations across human diseases and traits”. In: Nature Genetics 47.11 (Nov. 2015), pp. 1236–1241. ISSN: 1546-1718. DOI: 10.1038/ng.3406. URL: https://doi.org/10.1038/ng.3406.

[69] The Neale Lab. 2019. URL: https://ukbb-rg.hail.is/.

[70] Wei Zhou et al. “Efficiently controlling for case-control imbalance and sample relatedness in large-scale genetic association studies”. In: Nature Genetics 50.9 (Sept. 2018), pp. 1335–1341. ISSN: 1546-1718. DOI: 10.1038/s41588-018-0184-y. URL: https://doi.org/10.1038/s41588-018-0184-y.

[71] Matthew P. Conomos et al. “Model-free Estimation of Recent Genetic Relatedness”. In: The American Journal of Human Genetics 98.1 (2016), pp. 127–148. ISSN: 0002-9297. DOI: https://doi.org/10.1016/j.ajhg.2015.11.022. URL: https://www.sciencedirect.com/science/article/pii/S0002929715004930.

