## Supplementary Material for "Optimal strategies for learning multi-ancestry polygenic scores vary across traits"

Table 1: Relative variance of polygenic scores across traits and minority ancestry groups. Here, $\hat{g}_{EUR}^{min}$ denotes the vector of European-ancestry polygenic scores ($\mathrm{PGS_{EUR}}$) for the minority-ancestry test set, while $\hat{g}_{min}^{EUR}$ denotes the vector of minority-ancestry polygenic scores ($\mathrm{PGS_{min}}$) for the European-ancestry test set. Then, $\widetilde{\mathrm{Var}}(\hat{g}_{EUR}^{min})$ denotes the sample variance of $\hat{g}_{EUR}^{min}$ divided by the sample variance of $\hat{g}_{EUR}^{EUR}$.

| Trait | Minority ancestry | $\widetilde{\mathrm{Var}}(\hat{g}_{EUR}^{min})$ | $\widetilde{\mathrm{Var}}(\hat{g}_{min}^{min})$ | $\widetilde{\mathrm{Var}}(\hat{g}_{min}^{EUR})$ |
|---|---|---|---|---|
| Body mass index (BMI) | AFR | 0.582 | 0.201 | 0.152 |
| Mean corpuscular volume (MCV) | AFR | 0.573 | 1.768 | 0.346 |
| Mean corpuscular volume (MCV) | AMR | 0.941 | 1.124 | 1.084 |
| Mean corpuscular volume (MCV) | CSA | 0.870 | 0.360 | 0.337 |
| Mean corpuscular volume (MCV) | EAS | 0.837 | 1.239 | 0.673 |
| Mean corpuscular volume (MCV) | MID | 0.887 | 0.782 | 0.746 |
| Erythrocyte distribution width | AFR | 0.552 | 2.495 | 0.865 |
| Platelet crit | AFR | 0.548 | 0.383 | 0.212 |
| Platelet crit | AMR | 0.891 | 5.221 | 1.381 |
| Platelet crit | CSA | 0.807 | 0.366 | 0.271 |
| Platelet crit | EAS | 0.673 | 0.746 | 0.337 |
| Platelet crit | MID | 0.796 | 2.129 | 0.920 |
| Mean platelet volume | AFR | 0.543 | 0.089 | 0.068 |
| Lymphocyte count | AFR | 0.619 | 0.539 | 0.404 |
| Monocyte count | AFR | 0.605 | 0.365 | 0.269 |
| Eosinophill percentage | AFR | 0.616 | 0.257 | 0.196 |
| High light scatter reticuloctye count | AFR | 0.677 | 0.467 | 0.159 |
| Height | AFR | 0.558 | 0.050 | 0.039 |
| Height | AMR | 0.862 | 0.571 | 0.555 |
| Height | CSA | 0.780 | 0.059 | 0.053 |
| Height | EAS | 0.634 | 0.115 | 0.092 |
| Height | MID | 0.838 | 0.659 | 0.652 |
| Atrial fibrillation (AFib) | AFR | 0.766 | 22.395 | 16.436 |
| Diverticular disease of the intestine | AFR | 0.679 | 4.692 | 3.602 |
| Female genital prolapse | AFR | 0.654 | 220.417 | 79.755 |
| Female genital prolapse | CSA | 0.914 | 59.054 | 38.690 |
| Asthma | AFR | 0.614 | 3.414 | 2.531 |
| Asthma | AMR | 0.945 | 32.507 | 30.283 |
| Asthma | CSA | 0.903 | 10.497 | 10.240 |
| Asthma | EAS | 0.731 | 5.985 | 4.940 |
| Asthma | MID | 0.918 | 10.038 | 10.279 |
| Hypothyroidism | AFR | 0.611 | 2.707 | 2.077 |

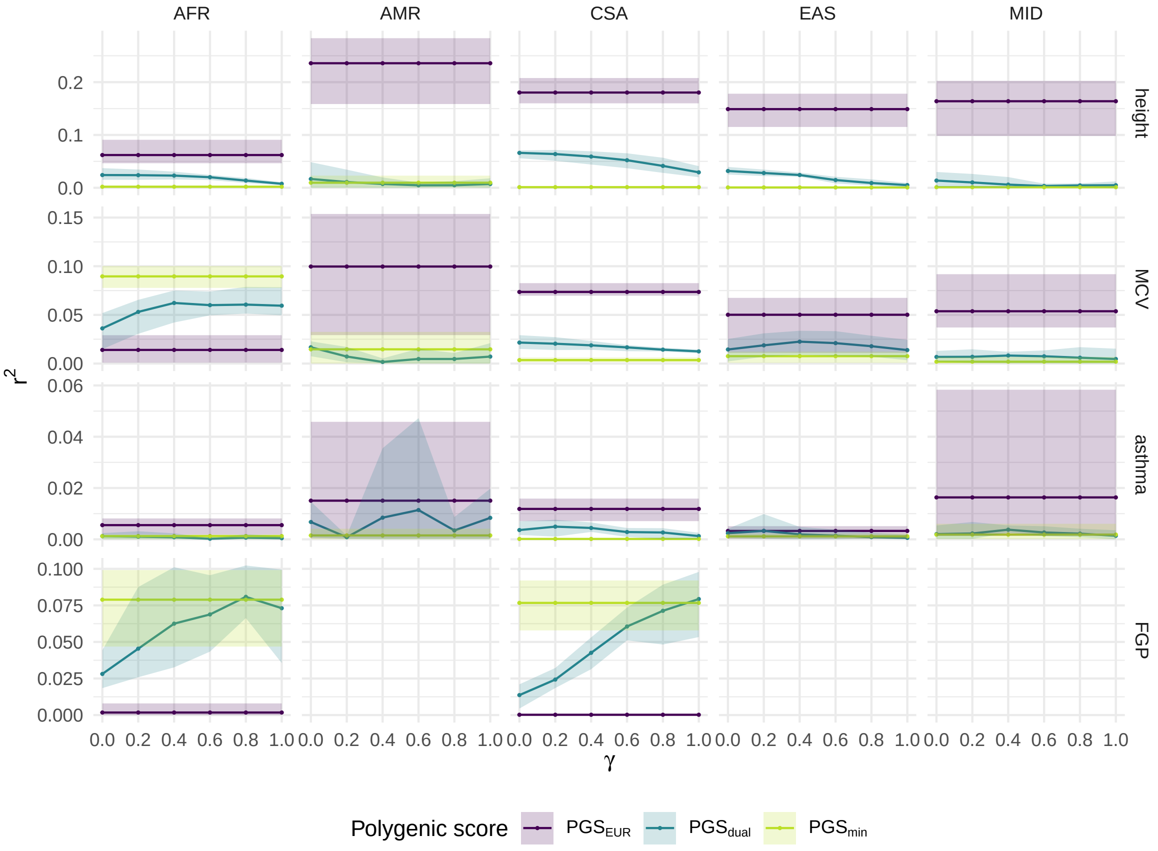


Figure 1: **Partial $r^2$ for European-, multi-, and minority-ancestry PGS on four traits in UK Biobank for five minority-ancestry groups using genotyped SNPs only.** The single-ancestry scores were estimated using a standard, unweighted LASSO. The dual-ancestry scores were constructed using an importance weighted LASSO with various degrees of reweighting $\gamma$. Error bars correspond to the range across five cross-validation rounds. The four traits considered are height, mean corpuscular volume (MCV), asthma, and female genital prolapse (FGP). We used inferred genetic ancestry labels from Pan-UKBB, with participants divided into six groups: European ancestry (EUR), African ancestry (AFR), Admixed American ancestry (AMR), Central/South Asian ancestry (CSA), East Asian ancestry (EAS), and Middle Eastern ancestry (MID). Analyses were not run on FGP for AMR, EAS and MID ancestry groups as the number of cases was fewer than 50 in each group.

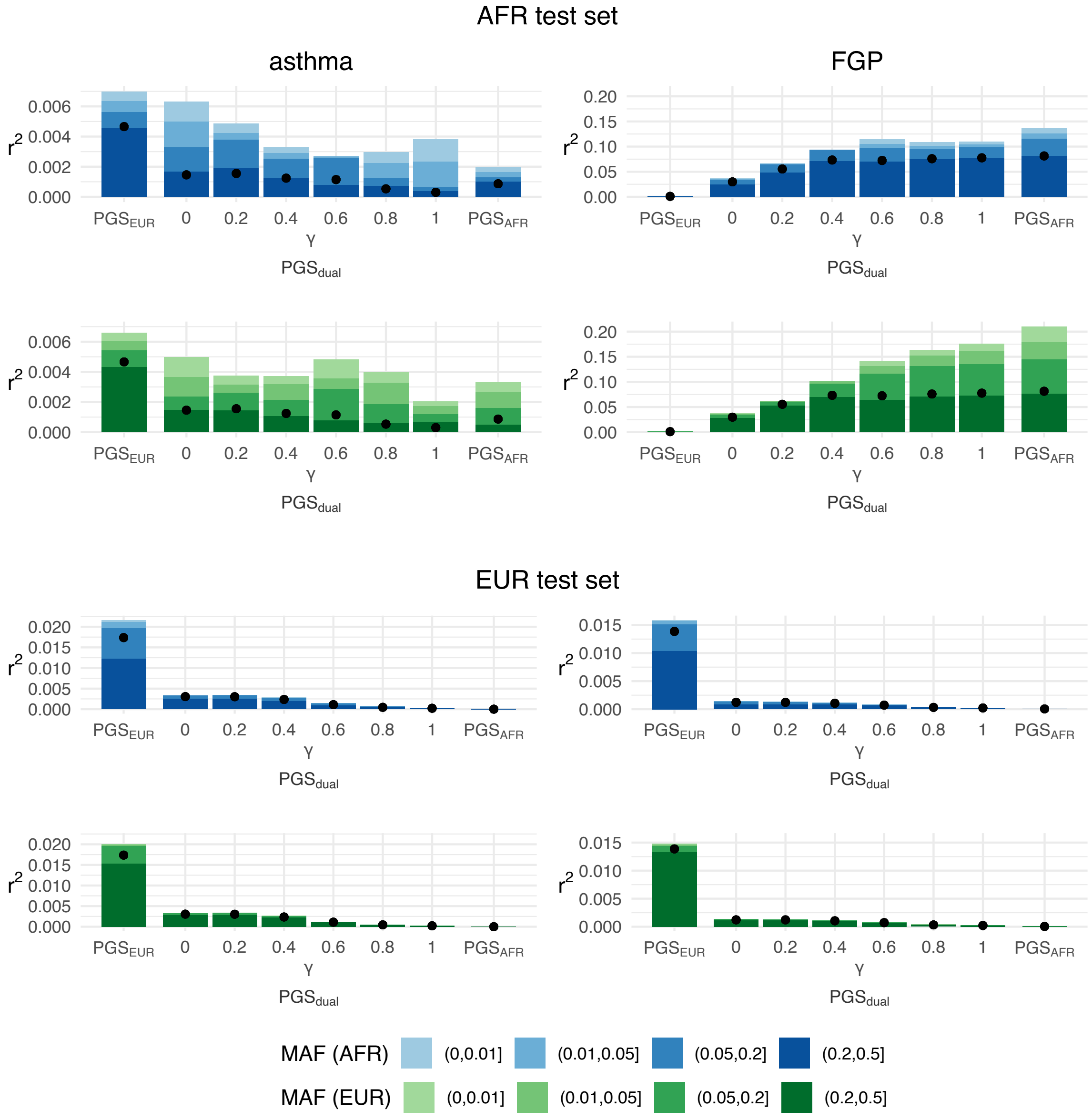


Figure 2: Allele frequency composition of variance explained by PGS for asthma (left) and female genital prolapse (right) in a African-ancestry test set (top) and a European-ancestry test set (bottom). The black dots represent partial $r^2$ for all the variants, i.e. the entire polygenic score. Variants were grouped according to their minor allele frequency in African-ancestry individuals (blue palette) or in European-ancestry individuals (green palette). Each bar represents the sum of the partial $r^2$ values for each subset of variants in a given polygenic score. Note that the height of the bar is generally higher than corresponding dot due to linkage disequilibrium between variants.

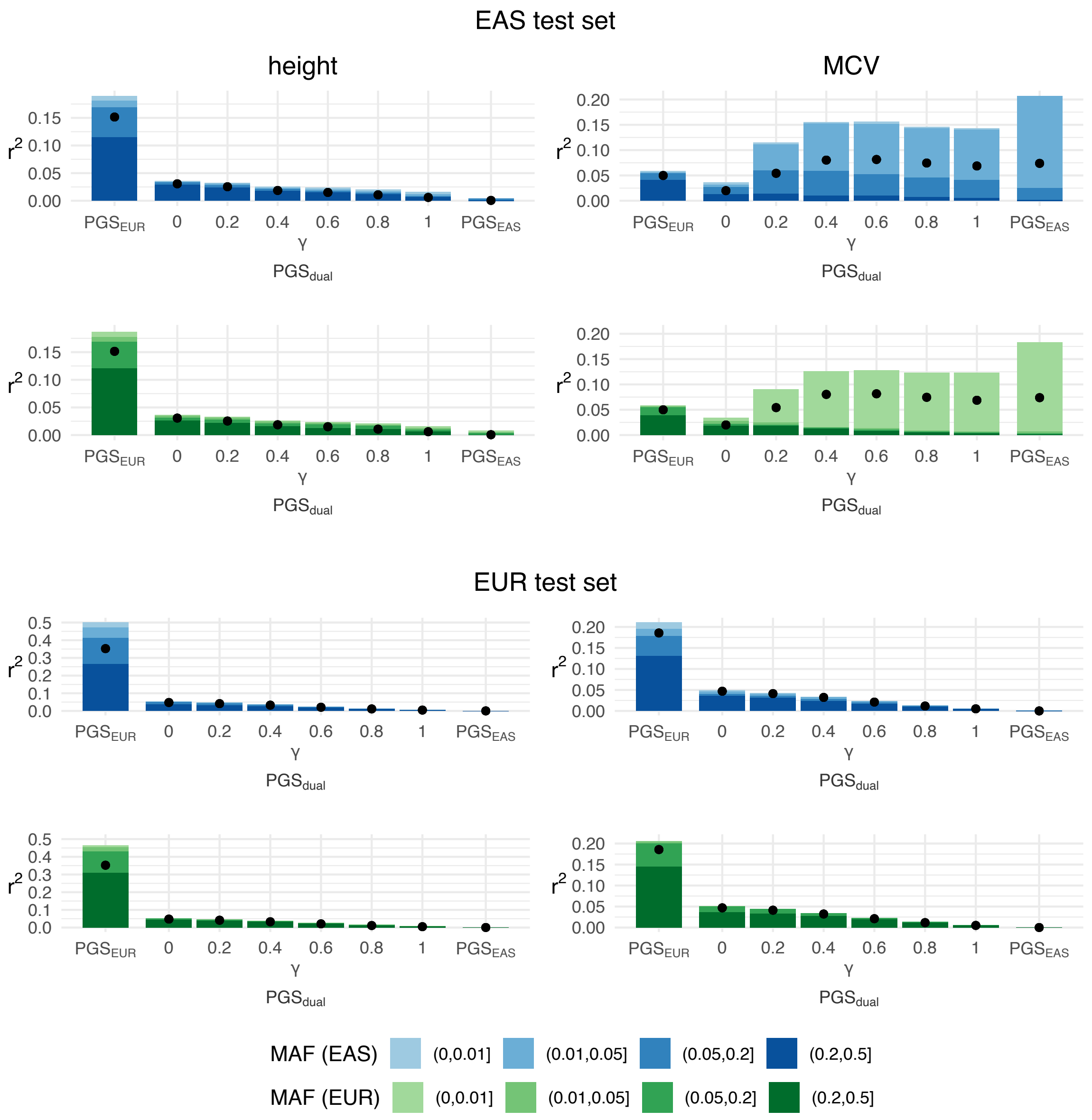


Figure 3: Allele frequency composition of variance explained by PGS for height (left) and mean corpuscular volume (right) in an East Asian ancestry test set (top) and a European-ancestry test set (bottom). The black dots represent partial $r^2$ for all the variants, i.e. the entire polygenic score. Variants were grouped according to their minor allele frequency in an East Asian ancestry individuals (blue palette) or in European-ancestry individuals (green palette). Each bar represents the sum of the partial $r^2$ values for each subset of variants in a given polygenic score. Note that the height of the bar is generally higher than corresponding dot due to linkage disequilibrium between variants.

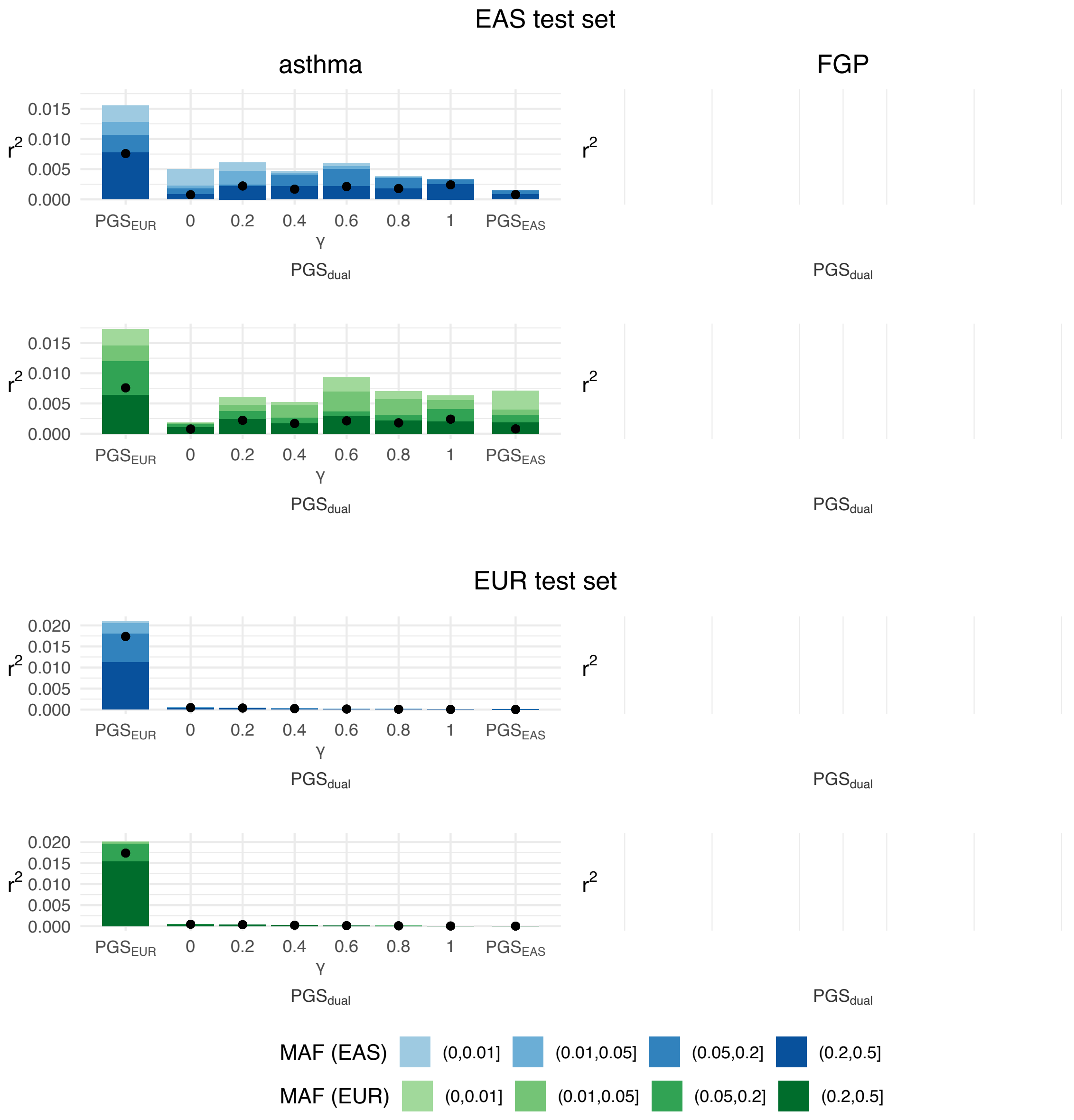


Figure 4: Allele frequency composition of variance explained by PGS for asthma (left) in an East Asian ancestry test set (top) and a European-ancestry test set (bottom). Analyses were not run on female genital prolapse as the number of cases was fewer than 50. The black dots represent partial $r^2$ for all the variants, i.e. the entire polygenic score. Variants were grouped according to their minor allele frequency in an East Asian ancestry individuals (blue palette) or in European-ancestry individuals (green palette). Each bar represents the sum of the partial $r^2$ values for each subset of variants in a given polygenic score. Note that the height of the bar is generally higher than corresponding dot due to linkage disequilibrium between variants.

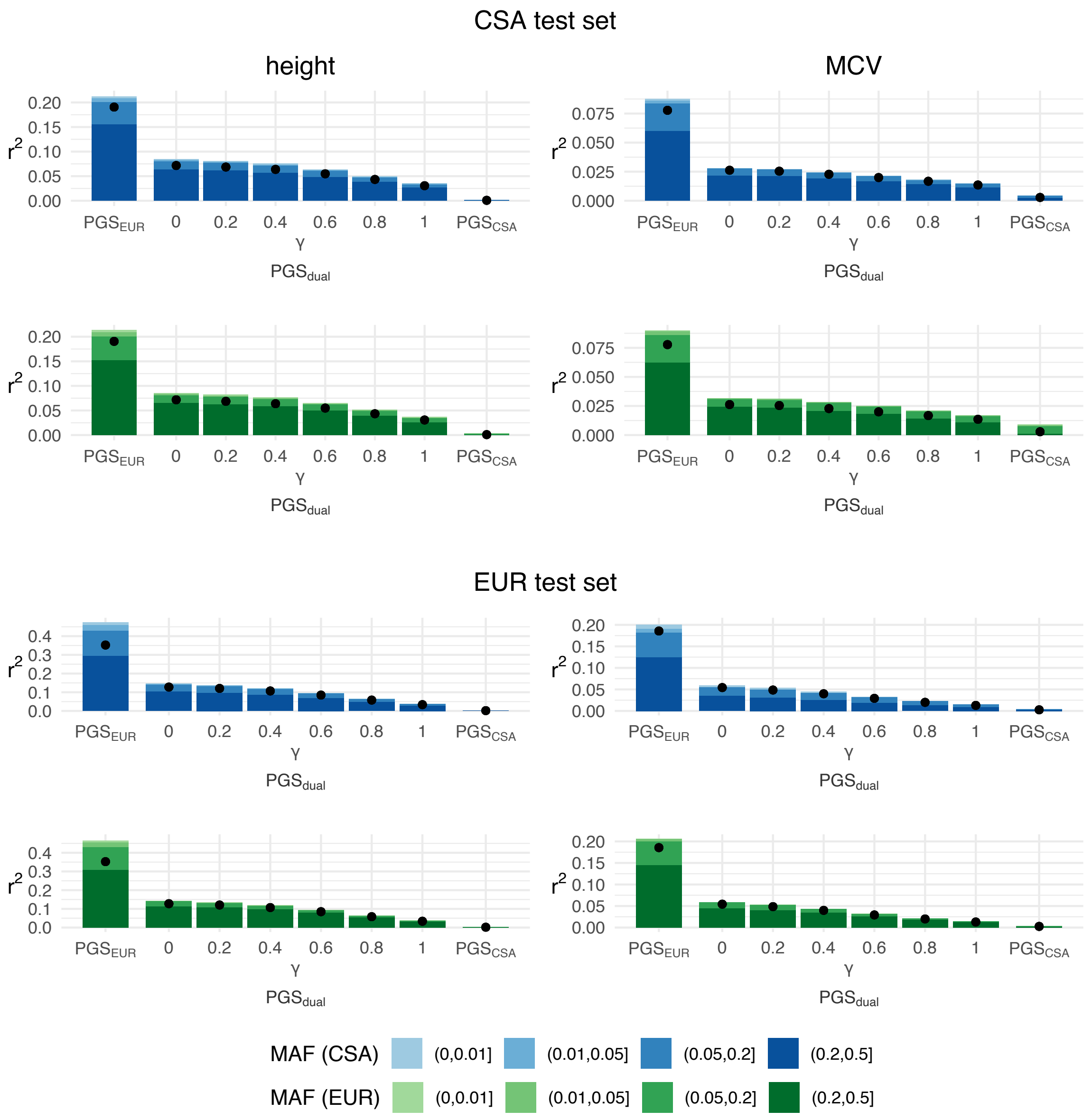


Figure 5: Allele frequency composition of variance explained by PGS for height (left) and mean corpuscular volume (right) in a Central/South Asian ancestry test set (top) and a European-ancestry test set (bottom). The black dots represent partial $r^2$ for all the variants, i.e. the entire polygenic score. Variants were grouped according to their minor allele frequency in a Central/South Asian ancestry individuals (blue palette) or in European-ancestry individuals (green palette). Each bar represents the sum of the partial $r^2$ values for each subset of variants in a given polygenic score. Note that the height of the bar is generally higher than corresponding dot due to linkage disequilibrium between variants.

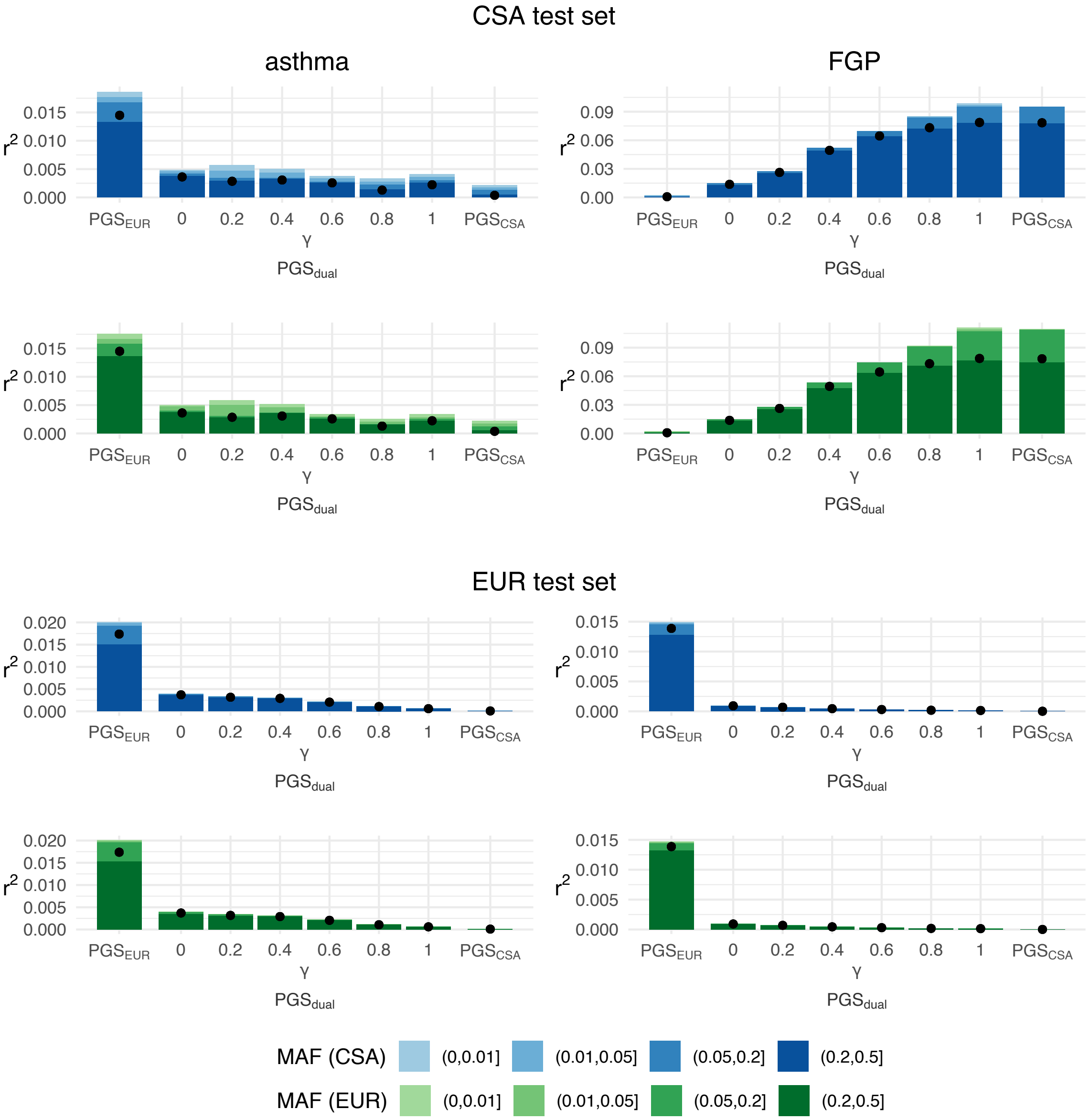


Figure 6: Allele frequency composition of variance explained by PGS for asthma (left) and female genital prolapse (right) in a Central/South Asian ancestry test set (top) and a European-ancestry test set (bottom). The black dots represent partial $r^2$ for all the variants, i.e. the entire polygenic score. Variants were grouped according to their minor allele frequency in a Central/South Asian ancestry individuals (blue palette) or in European-ancestry individuals (green palette). Each bar represents the sum of the partial $r^2$ values for each subset of variants in a given polygenic score. Note that the height of the bar is generally higher than corresponding dot due to linkage disequilibrium between variants.

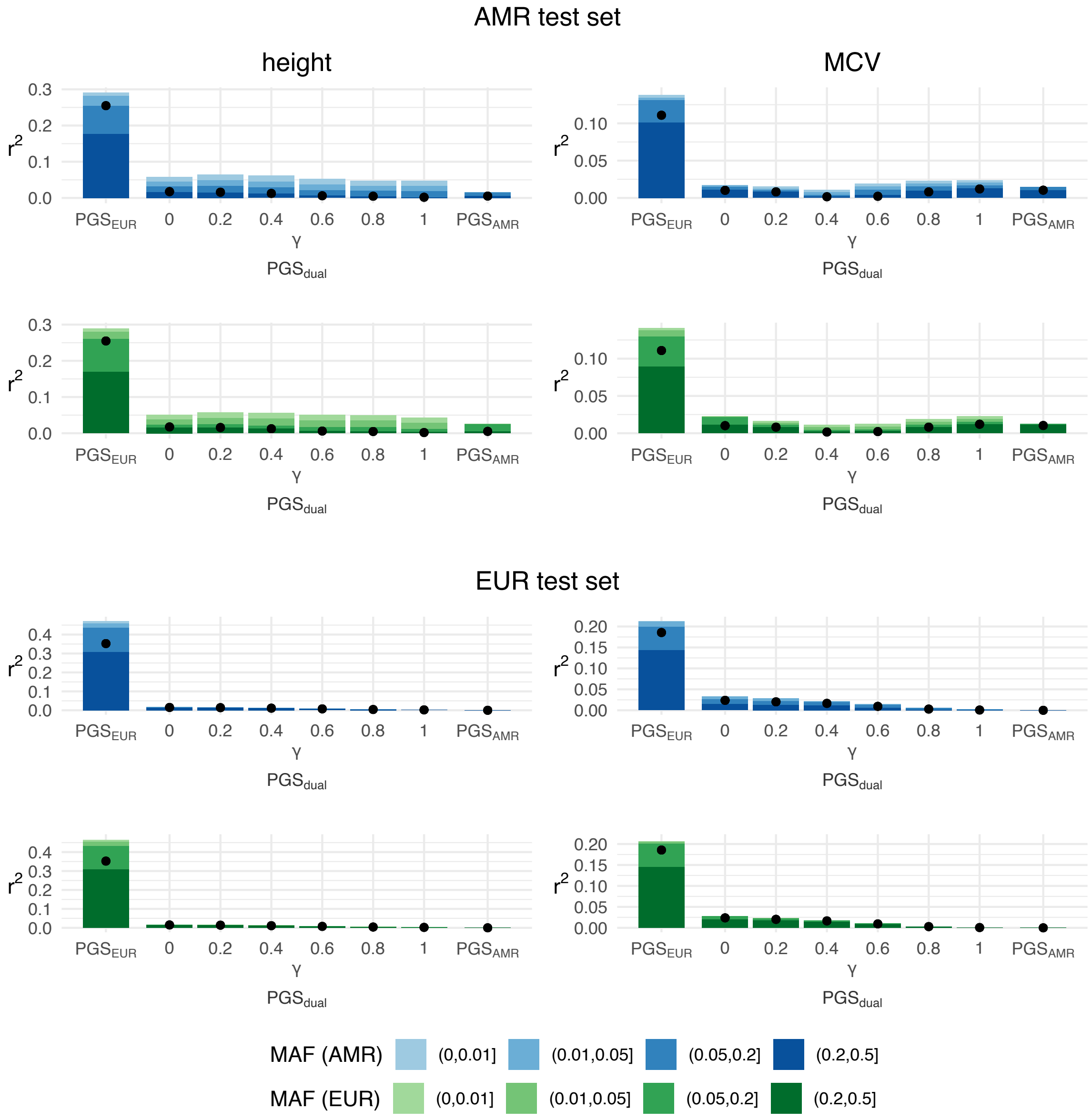


Figure 7: Allele frequency composition of variance explained by PGS for height (left) and mean corpuscular volume (right) in an Admixed American ancestry test set (top) and a European-ancestry test set (bottom). The black dots represent partial $r^2$ for all the variants, i.e. the entire polygenic score. Variants were grouped according to their minor allele frequency in an Admixed American ancestry individuals (blue palette) or in European-ancestry individuals (green palette). Each bar represents the sum of the partial $r^2$ values for each subset of variants in a given polygenic score. Note that the height of the bar is generally higher than corresponding dot due to linkage disequilibrium between variants.

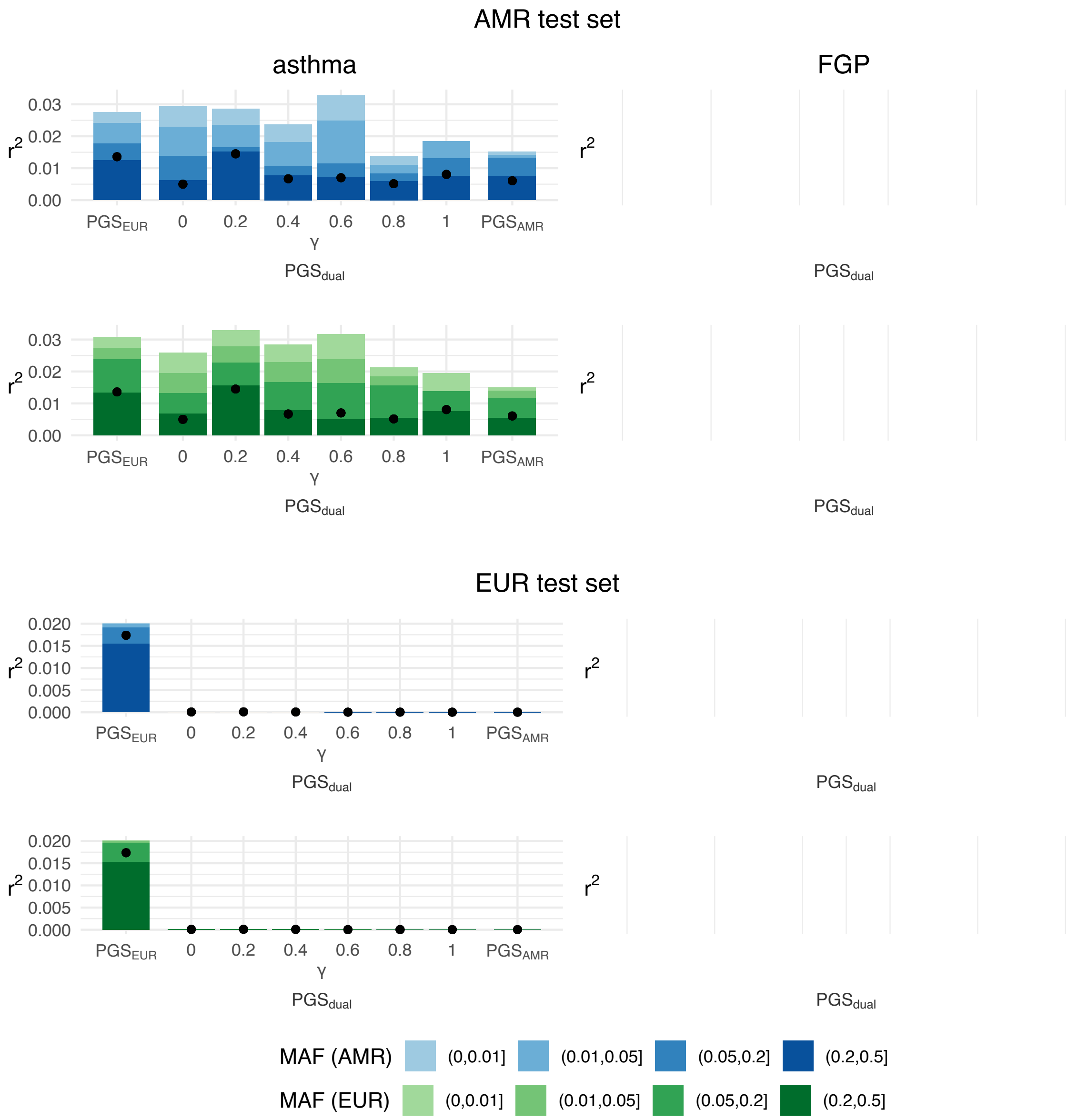


Figure 8: Allele frequency composition of variance explained by PGS for asthma (left) in an Admixed American ancestry test set (top) and a European-ancestry test set (bottom). Analyses were not run on female genital prolapse as the number of cases was fewer than 50. The black dots represent partial $r^2$ for all the variants, i.e. the entire polygenic score. Variants were grouped according to their minor allele frequency in an Admixed American ancestry individuals (blue palette) or in European-ancestry individuals (green palette). Each bar represents the sum of the partial $r^2$ values for each subset of variants in a given polygenic score. Note that the height of the bar is generally higher than corresponding dot due to linkage disequilibrium between variants.

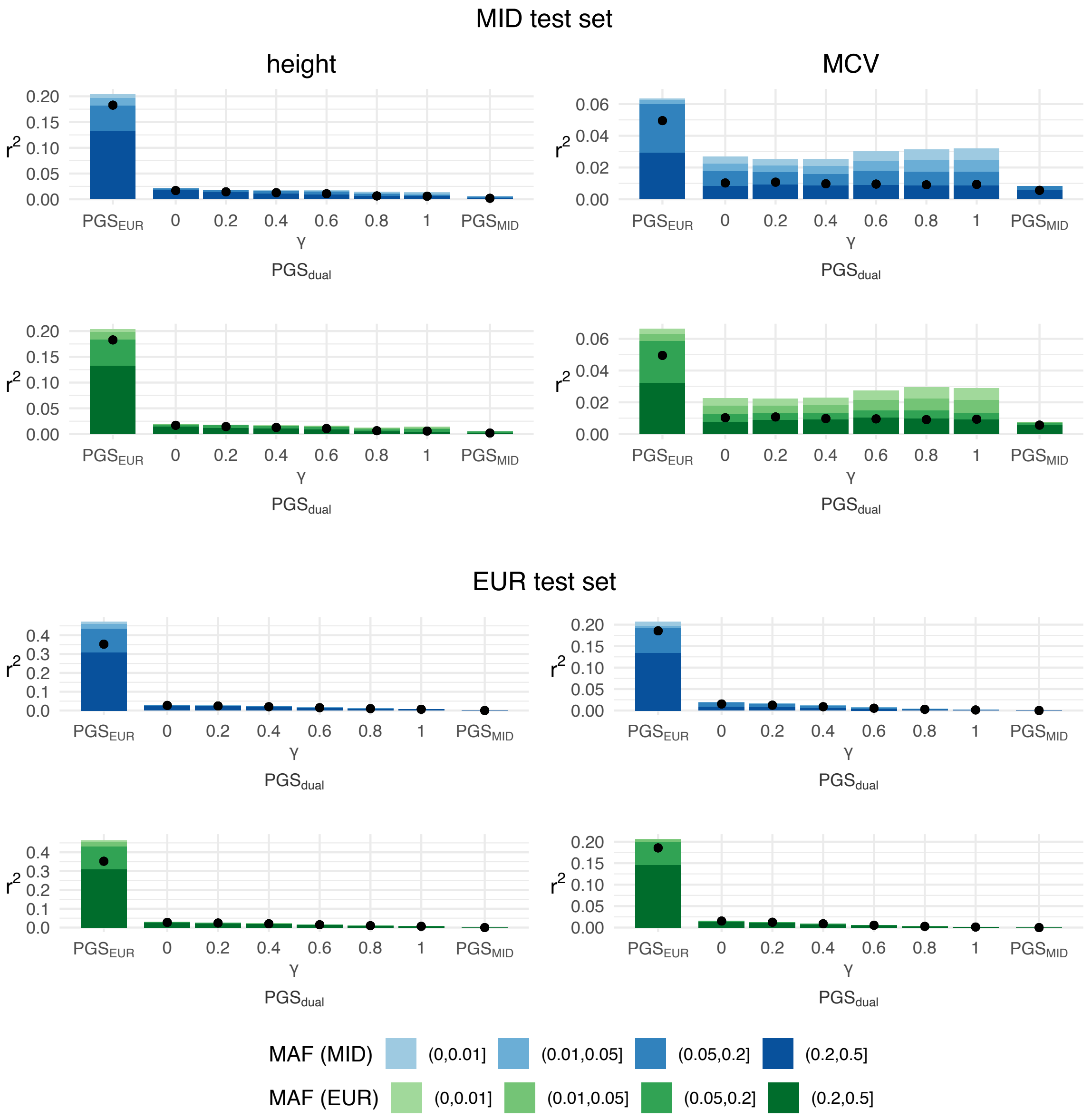


Figure 9: Allele frequency composition of variance explained by PGS for height (left) and mean corpuscular volume (right) in a Middle Eastern ancestry test set (top) and a European-ancestry test set (bottom). The black dots represent partial $r^2$ for all the variants, i.e. the entire polygenic score. Variants were grouped according to their minor allele frequency in a Middle Eastern ancestry individuals (blue palette) or in European-ancestry individuals (green palette). Each bar represents the sum of the partial $r^2$ values for each subset of variants in a given polygenic score. Note that the height of the bar is generally higher than corresponding dot due to linkage disequilibrium between variants.

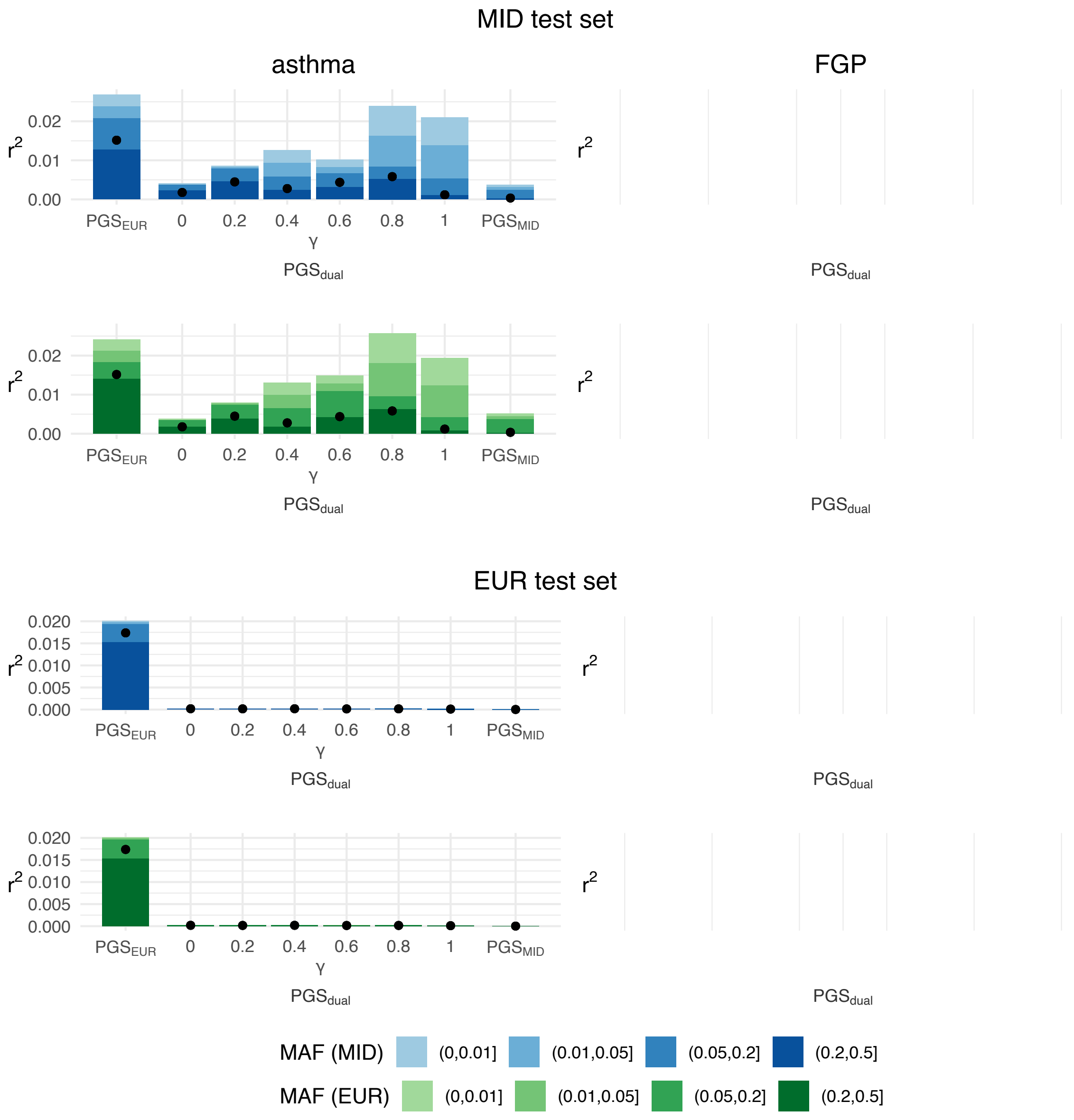


Figure 10: Allele frequency composition of variance explained by PGS for asthma (left) in a Middle Eastern ancestry test set (top) and a European-ancestry test set (bottom). Analyses were not run on female genital prolapse as the number of cases was fewer than 50. The black dots represent partial $r^2$ for all the variants, i.e. the entire polygenic score. Variants were grouped according to their minor allele frequency in a Middle Eastern ancestry individuals (blue palette) or in European-ancestry individuals (green palette). Each bar represents the sum of the partial $r^2$ values for each subset of variants in a given polygenic score. Note that the height of the bar is generally higher than corresponding dot due to linkage disequilibrium between variants.
